# Robust spatial hearing beyond primary interaural cues in humans over deep neural networks

**DOI:** 10.1101/2025.08.05.668776

**Authors:** Antonino Greco, Sangyeob Baek, Clara Rastelli, Markus Siegel, Christoph Braun

## Abstract

Spatial hearing allows humans to localize sound sources in the azimuth plane using interaural time (ITD) and level (ILD) differences, but the contribution of additional auditory features remains unclear. To investigate this, we measured human localization performance with natural and artificial stimuli that selectively included or excluded ITD and ILD as primary interaural cues. As expected, human listeners relied synergistically on ITD and ILD for accurate azimuth localization. Moreover, even when both primary cues were absent, localization performance remained above chance level. We compared human performance with state-of-the-art deep neural networks (DNN) optimized for sound localization to investigate possible computational mechanisms underlying this robust performance. In contrast to humans, DNNs demonstrated high accuracy only for stimuli that resembled their training regime but failed when primary interaural cues were absent. This human-DNN misalignment highlights a fundamental distinction in sensory processing strategies, potentially arising from the simplicity bias inherent in DNN training, with human reliance on a wider range of auditory features likely reflecting evolutionary pressures favoring adaptability across diverse acoustic environments. Together, our results demonstrate the robustness of human spatial hearing beyond primary interaural cues and point to promising directions for advancing artificial systems and informing clinical applications, such as cochlear implants and auditory prosthetics.

## Introduction

Sound localization is a fundamental auditory function that enables humans and other animals to navigate their environment, detect threats, and communicate effectively. Unlike vision, where spatial information is directly encoded by the retina and preserved in a topographically organized visual cortex, the auditory system does not have a direct spatial map at its sensory periphery. Instead, the brain infers the location of sounds through a complex integration of acoustic signals1. When sound waves interact with the unique shape of our outer ears, head, and upper torso, they are transformed in ways that produce distinctive, source-position dependent differences in timing, intensity, and spectral characteristics between the sensory inputs received by each ear^2,3^. It is well-known that in humans, binaural asymmetries in time and intensity, known as interaural time difference (ITD) and interaural level difference (ILD), serve as key spatial cues for localizing sound sources along the azimuth (horizontal) plane^4–6^. In addition, there is evidence showing that spectral cues are predominantly employed for elevation (vertical) localization and resolving front–back ambiguities rather than for determining azimuthal positions^2,7,8^. For instance, a study reported that human listeners, probed with artificial binaural sounds characterized by ITD, ILD and spectral cues, did not exploited monaural and interaural spectral cues to localize sounds on the azimuth plane^9^.

So far, most studies investigating the role of monaural and interaural cues in human spatial hearing have relied on artificially simple stimuli, such as pure tones or broadband white noise, to assess localization performance. Consequently, the acoustic features employed in sound localization within complex auditory scenes remain largely unexplored. Do humans prefer either ITD or ILD when faced with naturalistic stimuli or flexibly select the most appropriate depending on the sensory input? Do ITD and ILD suffice to predict human performance under naturalistic conditions? Furthermore, there is little evidence regarding human localization performance along the azimuth plane in the absence of the primary interaural cues, namely ITD and ILD. This raises the question of whether the auditory system may compensate for the loss of primary cues by relying on alternative acoustic features, or if the absence of primary cues inevitably leads to a loss of localization performance.

Recently, the advent of modern deep neural networks^10,11^ has opened new avenues for modeling sensory systems and optimizing them for real-world tasks^12–15^, achieving unprecedented levels of performance comparable to those of humans even on challenging sensory tasks^10,16–18^. This allows researchers to study a new kind of intelligent system, an “in silico” organism, which offers a unique opportunity to systematically investigate computational strategies underlying sensory processing and to compare them with biological organisms^19–28^. Specifically in the context of spatial hearing, a recent study^29^ demonstrated that many psychophysical characteristics of human sound localization can be ascribed to acoustic challenges of real-world environments, by training deep neural networks on realistic acoustic conditions and similar biological constrains that humans confront. These models successfully replicated several acoustic phenomena observed in humans, such as the frequency-specific reliance on ITD and ILD^9^, a bias for sound onsets^30^, and reduced localization ability when faced with simultaneous sound sources^31^. However, when trained in artificial environments devoid of natural elements like reverberation, ambient noise, and realistic soundscapes, model performance diverged markedly from human performance, illustrating how deep neural networks can be employed to uncover the real-world constraints governing sensory processing^29^. However, it is still unclear how these models perform under naturalistic conditions or when sensory inputs substantially diverge from their training regime, such as in the absence of primary interaural cues. Thus, investigating the acoustic features that drive localization performance of these models and how they relate to human preferred features may provide critical insights into the cognitive mechanisms of human spatial hearing.

To address these questions, we recorded a binaural spatial sound dataset using binaural recording microphones placed inside the ears of a human subject and naturalistic acoustic stimuli. We complemented these natural stimuli with artificially generated counterparts, which selectively included or excluded primary interaural cues, and tested humans and deep neural networks on these stimuli. This allowed us to examine the specific contribution of ITD, ILD and other spatial features to the localization performance under both natural and out-of-distribution conditions. Notably, some artificially generated stimuli completely lacked the primary interaural cues, which allowed us to test the robustness of the human and model sensory processing and reveal the impact of alternative acoustic features independently of ITD and ILD cues. We further investigated the nature of these alternative features to the primary interaural cues by additionally generated artificial stimuli with binaurally matched spectral components and evaluate the human and model responses to these stimuli.

## Results

### Benchmarking spatial hearing with natural and artificial sounds

We collected a dataset of spatial sounds to train (finetuning) deep neural networks (DNN) models for sound localization and to test both humans and DNNs. We recorded 50 natural sounds^32^, frequently heard in everyday life by humans, using binaural recording microphones placed inside the ears of a human (Fig. 1A). This setup allowed us to study the contribution of ITD and ILD to sound localization performance (Fig. 1B). We played source sounds from a speaker located at 9 different positions on the azimuth plane with respect to the human listener and the binaural recording system (Fig. 1C). Recordings were performed on 3 different acoustic environments (Fig. 1D), one anechoic room, one large lecture hall (indoor) and in a garden (outdoor). After collection of the spatial sound dataset, we split the data in one set for finetuning DNNs (Fig. S1A-B) and one for testing both humans and DNNs (Fig. 1A), based on the alignment of the ITD and ILD features to the azimuth position and semantic features extracted from the YAMNet^33,34^ model (Fig. S1C-D, see methods). As a sanity check, we plotted the ITD and ILD of the final sets (test set on Fig. 1E, train set on Fig. S1B) as a function of the azimuth position, confirming the linear relationship between the interaural cues and the azimuth angle.

**Figure 1.**
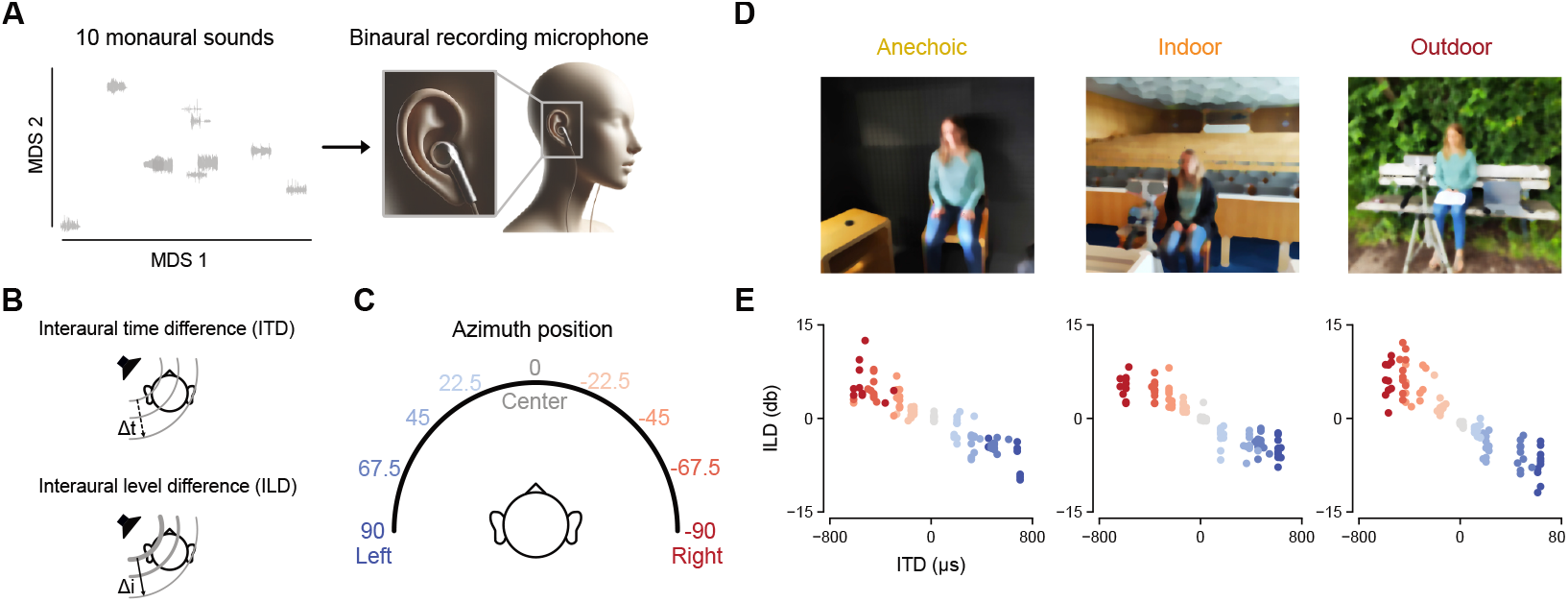
Binaural recording of the spatial sound dataset. **a**. We sampled 50 monaural sounds from the Natural Sound Stimulus Set dataset^32^ and used 10 of them to test humans and deep neural networks on spatial hearing. On the left, we projected these sounds on a 2D plane using multidimensional scaling (MDS) based on the distance matrix of the sound waveforms. On the right, pictorial illustration of the binaural recording settings we used to collect the spatialized sounds. This consisted of placing two microphones inside the ears of a human and playing the sounds from a speaker standing in 9 different azimuth locations at 0° elevation. **b**. Graphical illustration of the interaural cues (ITD and ILD) investigated in this study. **c**. Azimuth locations where we presented the monaural sounds and collected the binaural recording via the microphones inside the human’s ears. **d**. Photographs depicting the 3 environments where we recorded the spatialized sounds. **e**. Scatterplots showing the ITD and ILD of the resulted sounds across the 3 environments and color-coded as a function of the azimuth position.

Next, we tested human listeners using our natural binaural recordings alongside two sets of artificial stimuli, which we refer to as “synthesized” and “defeaturized” stimuli, to isolate ITD and ILD contributions (Fig. 2A). For synthesized stimuli, we took the monaural source sounds and imposed the exact ITD, ILD, or both (denoted ITD-ILD) measured from the outdoor recordings. For defeaturized stimuli, we removed ITD, ILD, or both from the outdoor binaural recordings, leaving only other “third” acoustic features. Thus, natural sounds contained ITD, ILD, and third features, synthesized sounds contained only ITD and/or ILD, and defeaturized sounds contained only third features.

**Figure 2.**
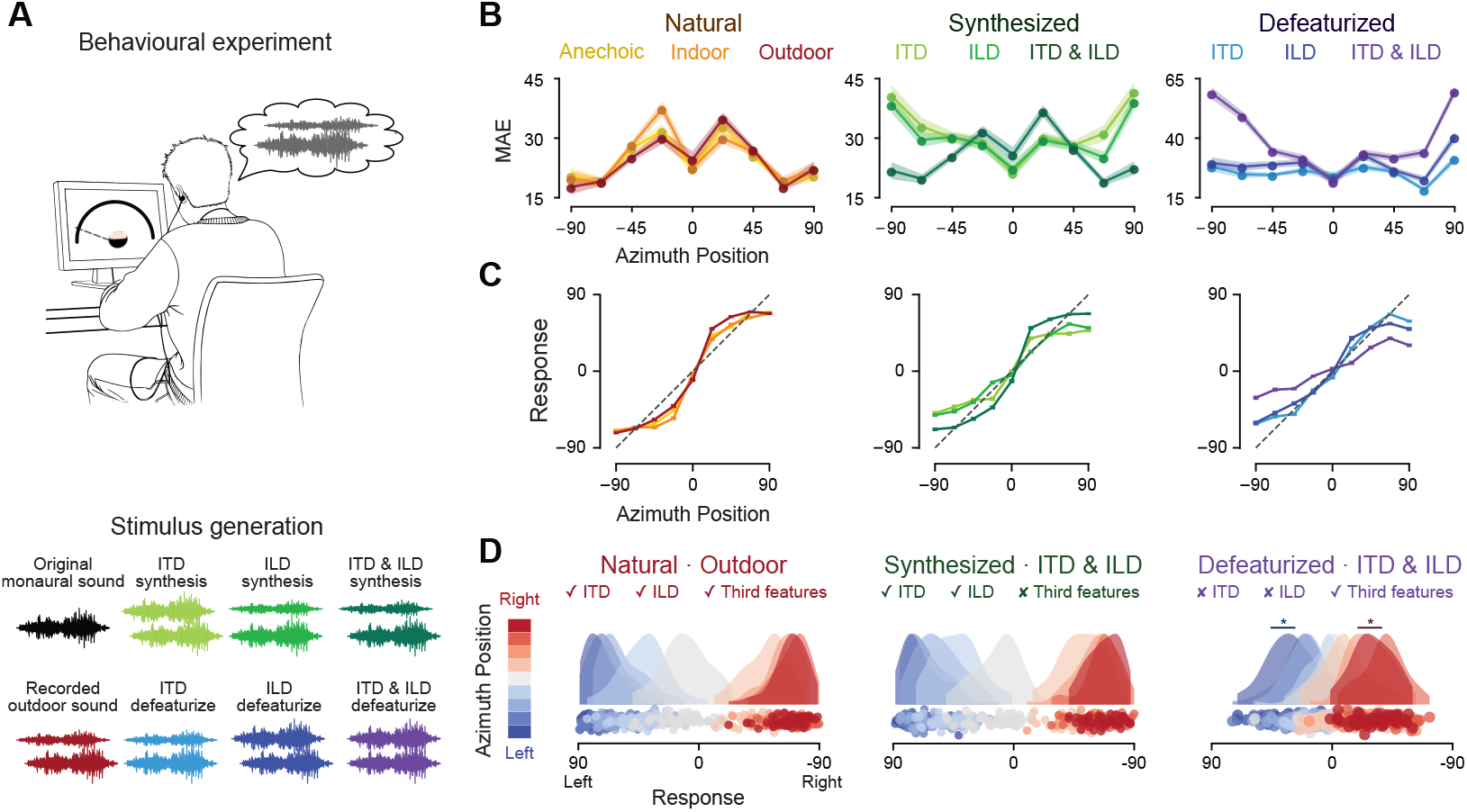
Human performance on natural and out-of-distribution sounds. **a**. We probed human listeners using our recorded spatial sounds and additional artificial stimuli generated to investigate the contribution of ITD and ILD to their sound localization performance. Top, graphical illustration of the experimental setup we used to deliver the stimuli and collect the response. Bottom, schematic illustration of the stimulus generation procedure. In the synthesis stimuli, the original monaural sound is binaurally either shifted in time (ITD) or amplified (ILD) or both (ITD & ILD). In the defeaturize sounds, the recorded outdoor sound has the binaural time shifts (ITD) or amplifications (ILD) or both (ITD & ILD) removed. **b**. Lineplots depicting the mean absolute error (MAE) in degrees between the true azimuth and the human responses as a function of the azimuth position across stimulus types. Shaded areas represent SEM. **c**. Lineplots showing the transfer function of the group-average human performance mapping the true azimuth position (x-axis) to the responded one (y-axis). Dotted black lines indicate ideal pattern, while horizontal bars indicate SEM. **d**. Response distributions for the natural outdoor, synthesis ITD-ILD and defeaturize ITD-ILD stimuli. Each dot represents the average response of each participant to a specific location, color-coded by the azimuth position as expressed by the colorbar. Asterisks represent statistical significance.

### Characterizing human sound localization performance

We first investigated reaction times (RT) and confidence scores of human participants. There was no significant difference between RTs across stimulus types and azimuth position (Fig. S2A; all P^FDR^ > 0.05). Across all stimulus types, confidence scores were descriptively higher in lateral compared to central positions (Fig. S2B) and significantly correlated (Fig. S2C) with absolute ITD (r = 0.129, P < 0.0001) and ILD (r = 0.185, P < 0.0001). Thus, human participants were more confident for stronger interaural cues.

Next, we investigated localization errors by plotting the mean absolute error (MAE, Fig. 2B) and accuracy (Fig. S3A) as a function of azimuth position. Qualitatively, the performance for synthesized ITD-ILD stimuli resembled natural performance, exhibiting an M-shaped MAE pattern across azimuth positions (Fig. 2B). As expected, removing primary interaural cues (defeaturized ITD-ILD stimuli) degraded performance, with highest error rates at lateral positions. We computed the distance matrix between all stimulus types by comparing their error pattern across azimuth positions (Fig. S3B). This revealed three clusters of performance. The first cluster concerned natural and synthesized ITD-ILD stimuli, the second the synthesized and defeaturized with either ITD or ILD and the third only included defeaturized ITD-ILD stimuli.

### Human spatial hearing is robust to the absence of ITD and ILD

We next directly quantified the mapping between true and reported azimuths (Figs. 2C and Fig. 2D; see Fig. S3C for 2D-density plots). We found that both natural and synthesized ITD-ILD stimuli produced response distributions concentrated at the center and extreme lateral positions (Fig 2C). Defeaturized ITD-ILD stimuli showed a different pattern (Fig. 2C right), but the transfer function for these stimuli was not entirely flat, suggesting that for these stimuli without ITD or ILD participants used remaining third features for sound localization. Also visualizing response distributions as a function of azimuth position (Fig. 2D) showed that, for defeaturized ITD-ILD stimuli, the overall response patterns remained remarkably aligned with the actual sound source locations (Fig. 2D right), albeit more compressed toward the center compared to the natural and synthesis ITD-ILD (Fig 2D left and middle; for all the other stimulus types, see Fig. S3D). This effect was statistically significant. Humans significantly localized defeaturized ITD-ILD stimuli correctly to the left (P_FDR_ < 0.0001, d = 2.01, 95% CI [1.52, 2.5]) or right (P_FDR_ < 0.0001, d = 1.73, 95% CI [1.29, 2.18]). Thus, third non-ITD/ILD features alone provided enough information for basic lateralization of human listeners.

### Humans rely on third features for localization performance

Next, we quantitatively investigated how much spatial cues contributed to human localization performance for different stimuli, by computing a metric of central and lateral performance using a weighted average of the MAE across the azimuth positions (Fig. 3A-B). First, we tested all combinations (anechoic-indoor, indoor-outdoor, anechoic-outdoor) of environments in the recorded natural sounds. We found no difference of centralized and lateralized MAE for any of the environment combinations (all P_FDR_ > 0.05), suggesting that humans were not affected by the environment under natural conditions. Then, we compared natural outdoor sounds with the synthesized and defeaturized ITD-ILD stimuli, since the latter were generated from the outdoor environment statistics. No difference was observed for centralized MAE (all P_FDR_ > 0.05), while lateralized MAE was significantly lower for natural stimuli compared to synthesized (P_FDR_ = 0.048, d = 0.31, 95% CI [0.02, 0.6]) and defeaturized (P_FDR_ < 0.0001, d = 3.43, 95% CI [2.69, 4.17]) ITD-ILD stimuli. MAE was significantly lower for synthesized ITD-ILD as compared to defeaturized ITD-ILD stimuli for lateral positions (P_FDR_ < 0.0001, d = 3.09, 95% CI [2.41, 3.76]) but not for central positions (P_FDR_ > 0.05). These results were confirmed by averaging the MAE across all azimuth positions (Fig. S4A, all P_FDR_ < 0.003). This suggests that the lack of both ITD and ILD strongly impacts human performance. However, for lateral stimuli ITD and ILD presence alone in synthesized stimuli does not enable localization performance as for natural stimuli.

**Figure 3.**
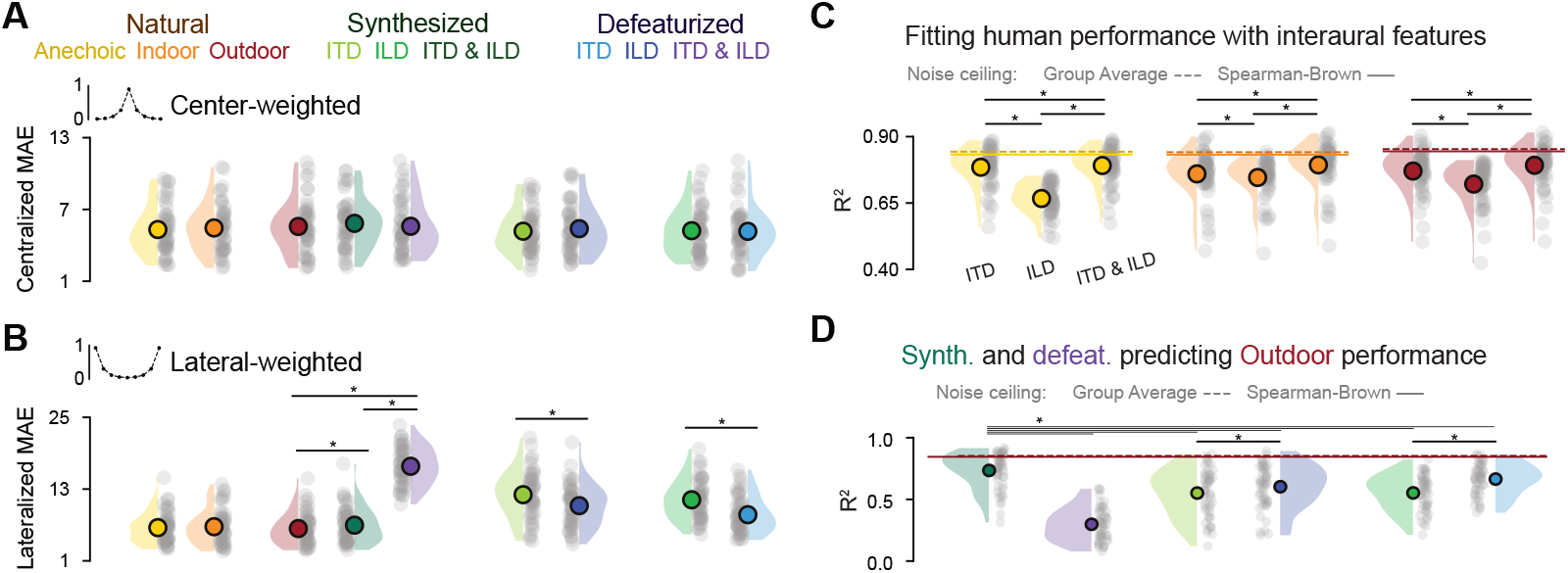
Human spatial hearing within and beyond ITD and ILD. **a**. Raincloud plots showing the centralized MAE score in degrees using the center-weighted kernel showed on top (y-axis of the kernel plot indicating the weight values). **b**. Raincloud plots showing the lateralized MAE score in degrees using the lateral-weighted kernel showed on top. Asterisks represent statistical significance. **c**. We tested the specific contribution of ITD and ILD to the human localization performance under natural settings. Raincloud plots show R^2^ scores for linear regression models fitted to predict human responses on each natural environment and using as regressor either the ITD, the ILD or both computed on the natural sounds. Each dot represents a participant, while the circle overlapping the density plot represent the mean. Horizontal lines represent noise ceiling estimates, either from the group average (dotted) or with the Spearman-Brown method (solid). Asterisks signal statistical significance, generally as P < 0.05. **d**. We used human responses to synthesis and defeaturize stimuli to predict responses to natural outdoor stimuli. Raincloud plots show R^2^ scores for each condition, with each dot representing a single participant and circles representing the group-level mean. Horizontal lines represent noise ceiling estimates on the natural outdoor environment (hence, the red color) and asterisks depict statistical significance.

Our stimulus design allowed us to directly test the contribution of third features. Specifically, we compared stimuli that had identical ITD or ILD but differed only in the inclusion (defeaturize stimuli) or exclusion (synthesis stimuli) of the third features. In the first comparison, we matched ITD and compared synthesized ITD with defeaturized ILD stimuli. In the second comparison, we contrasted synthesized ILD with defeaturized ITD stimuli. Crucially, we found that lateralized MAE was lower in defeaturized as compared to synthesized stimuli (Fig. 3B), both when comparing ITD-matched (P_FDR_ < 0.0001, d = 0.67, 95% CI [0.36, 0.98]) and ILD-matched (P_FDR_ < 0.0001, d = 1.26, 95% CI [0.88, 1.64]) stimuli. There was no difference for centralized MAE (Fig. 3A, all P_FDR_ > 0.05). We confirmed these results for the average MAE across all azimuth positions (Fig. S4A, all P_FDR_ < 0.0001). These results provide direct evidence that humans localize sounds not only based on ITD and ILD, but also use third features.

### ITD and ILD alone do not fully explain natural human performance

We adopted a regression-based approach to quantitatively compare the unique contribution of ITD, ILD and third features. How much of the behavioral variance can be explained by ITD and ILD alone? To answer this, we fitted human responses using the primary interaural cues from the sounds they heard (Fig. 3C). Across all natural environments, ITD was a significantly stronger predictor of human responses than ILD (all P_FDR_ < 0.009, all Cohen’s d > 0.38). Furthermore, the combination of ITD and ILD provided a significantly better prediction than either ITD (R^2^: all P_FDR_ < 0.0001, all d > 0.72; adjusted-R^2^: all P_FDR_ < 0.002, all d > 0.46) or ILD (R^2^: all P_FDR_ < 0.0001, all d > 2.23; adjusted-R^2^: all P_FDR_ < 0.0001, all d > 1.90) alone (see Fig. S4B for artificial*-s*timuli and Fig. S4C for MAE*-b*ased regression).

Crucially, we also tested whether primary interaural cues explained all the variance of human responses by computing a noise-corrected version of R^2^. As listeners retained some localization ability even without primary interaural cues, we predicted that ITD and ILD would not fully account for human behavior under natural listening. Indeed, for all natural environments, we found that the combined ITD and ILD model explained significantly less than 100 % of the behavioral variance (noise-corrected R^2^: 94.0%, 94.5%, and 92.8% for anechoic, indoor, and outdoor environments, respectively; R^2^ < 1: all P_FDR_ < 0.0001, all d > 1.44). Thus, even in natural environments a small but significant portion of localization performance depended on additional cues beyond ITD and ILD.

### Human spatial hearing beyond ITD and ILD

We next investigated the predictive power of human responses to artificial stimuli that specifically manipulated ITD and ILD cues. Instead of fitting ITD and ILD features to human responses, we utilized each participant responses to manipulated sounds to predict its performance in the naturalistic (outdoor) setting (Fig. 3D). As in the previous analysis, we focused on explained behavioral variance (see Fig. S4D for MAE based regression). We found that responses to synthesized ITD-ILD stimuli were the best predictor of localization performance in the outdoor environment (all P_FDR_ < 0.0001, all d > 0.78) with a noise-corrected R^2^ of 85.7%, while the worst was the defeaturize ITD-ILD, explaining only the 34.6% of the variance.

We also compared the predictability of human performance on natural stimuli of the synthesized ITD with the defeaturized ILD stimuli and the synthesized ILD with the defeaturized ITD stimuli. As above, this allowed us to directly compare the contribution of third features to predicting human performance on natural stimuli. Crucially, we found significantly higher R^2^ scores for defeaturized as compared to synthesized stimuli, both for ITD-matched (P_FDR_ = 0.005, d = 0.41, 95% CI [0.12, 0.7]) and ILD-matched (P_FDR_ < 0.0001, d = 1.28, 95% CI [0.9, 1.66]) stimuli. Again, this provides direct evidence that humans rely on third features beyond ITD and ILD.

### Characterizing deep neural network localization performance

We next evaluated the performance of deep neural network models trained for sound localization and compared them to human behavior. Specifically, we investigated the behavior of deep convolutional neural networks^10,11^ (CNN) pretrained in a previous study^29^, since these models, designed to operate under realistic acoustic conditions, have demonstrated high accuracy in localizing sounds while exhibiting perceptual characteristics closely aligned with human spatial hearing^29^. The model architecture comprises a hierarchy of convolutional layers, culminating in fully-connected layers that generate the final sound location prediction (Fig. 4A). The model receives a cochleagram of the binaural sound waveform as input, which is a biologically plausible representation of the instantaneous mean firing rates of the auditory nerve, reflecting the human cochlea’s output^35^. We evaluated the localization performance of the 10 top-performing CNN models, from the pretraining phase^29^, on both natural and out-of-distribution stimuli, treating them as individual participants analogously to human listeners.

**Figure 4.**
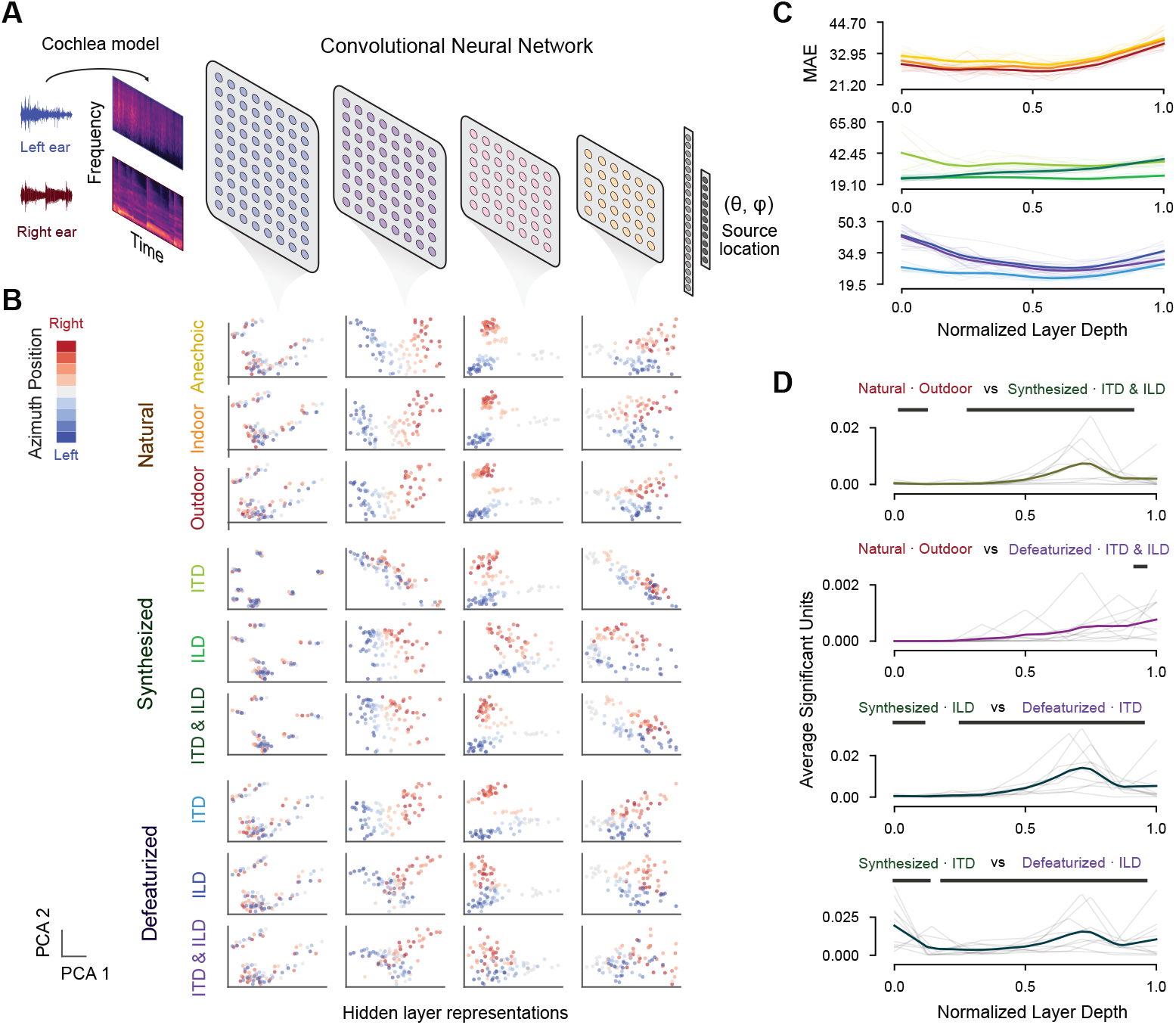
Probing deep neural networks optimized for sound localization on natural and out-of-distribution sounds. **a**. Schematic illustration of the model architecture of the Convolutional neural network (CNN) models we used. The input binaural audio signal was passed through a computational model of the human cochlea and the resulting time-frequency representation (cochleagram) was the input of the CNN model. Then, a series of convolutional blocks extracted relevant features for sound localization and the final fully-connected layers performed the prediction over the azimuth (θ) and elevation plane (*φ*). Below in **b**., scatterplots depicting the hidden layer representations of an exemplar CNN model, defined as the first two principal components of the layer activations across all stimuli. Each column shows a different layer across the hierarchy of the convolutional blocks, from left to right being early to late layers, while each row shows the representational space of different stimulus types. Each dot represent a single sound at each azimuth position and is color-coded according to the actual azimuth position. **c**. Lineplots showing the mean absolute error (MAE), resulting by predicting the azimuth position from the layer activations, as a function of the normalized layer depth. Lines are color-coded by the stimulus types with thin lines represent each CNN model, while thick lines show the average across all models **d**. Lineplots depicting the average number of significant units resulting from specific comparisons between stimulus types as a function of the normalized layer depth. Thin lines represent each CNN model, while thick lines show the average across all models. Horizontal bars indicate statistical significance.

First, we qualitatively inspected hidden layer representations of these models to investigate how they processed spatialized sounds (Fig. 4B). The natural stimuli’s representational geometry showed a clear separation between leftward and rightward azimuths, especially in middle layers, a pattern closely mirrored by synthesized ITD-ILD representations, whereas defeaturized stimuli, especially in absence of both primary interaural cues, showed markedly dispersed activations with weak azimuth clustering. To further investigate how these hidden representations relate to the spatial location of the input stimuli, we tried to predict the sound location from the layer activations (Fig. 4C).

Localization error was lowest in middle layers, indicating that spatial cues are progressively transformed and then condensed into abstract, decision*-l*evel representations in later layers.

Then, we adopted a “systems neuroscience” approach^36,37^ to study the feature selectivity of models units by comparing their activity between different types of stimuli (Fig. 4D). This allowed us to detect whether a unit or a layer was sensitive to some specific acoustic features. When contrasting the natural outdoor sounds with the synthesis ITD-ILD, we found that the number of units which differed in their activity between these sounds was significantly higher than zero in early (all P_FDR_ < 0.023, all d > 0.92) and middle-to-late layers (all P_FDR_ < 0.020, all d > 0.94). Differently, contrasting the natural related unit activations with the defeaturized ITD-ILD resulted in only late layers exhibiting a significant number of selective units (all P_FDR_ < 0.020, all d > 1.43). This suggests that primary interaural cues were encoded only in late stages of the model hierarchy.

Finally, we also compared pairs of stimulus sets that share identical ITD or ILD, similarly to the above analyses for humans. Again, we observed early (all P_FDR_ < 0.024, all d > 0.88) and middle-to-late (all P_FDR_ < 0.024, all d > 0.90) layers showing significantly selective units, suggesting that models were encoding third features in these layers.

### Deep neural networks are not robust to the absence of ITD and ILD

We next investigated CNN behavior using the same metrics that we used for human behavior. Models qualitatively reproduced the M-shaped MAE pattern found in humans (Fig. 5A), with lower MAE for lateral positions, which was similarly restored in synthesized ITD-ILD and synthesized ITD stimuli, again suggesting an ITD bias. Notably, we found a different error profile for all the other artificial stimulus types, with defeaturized ITD-ILD showing the highest error. Next, by inspecting their transfer functions (Fig. 5B), we observed that models closely followed the ideal diagonal for natural stimuli, especially in anechoic and indoor settings, and for synthesis ITD and ITD-ILD, but not for synthesis ILD, indicating reliance on ITD. The largest deviations occurred for defeaturized ITD-ILD stimuli.

**Figure 5.**
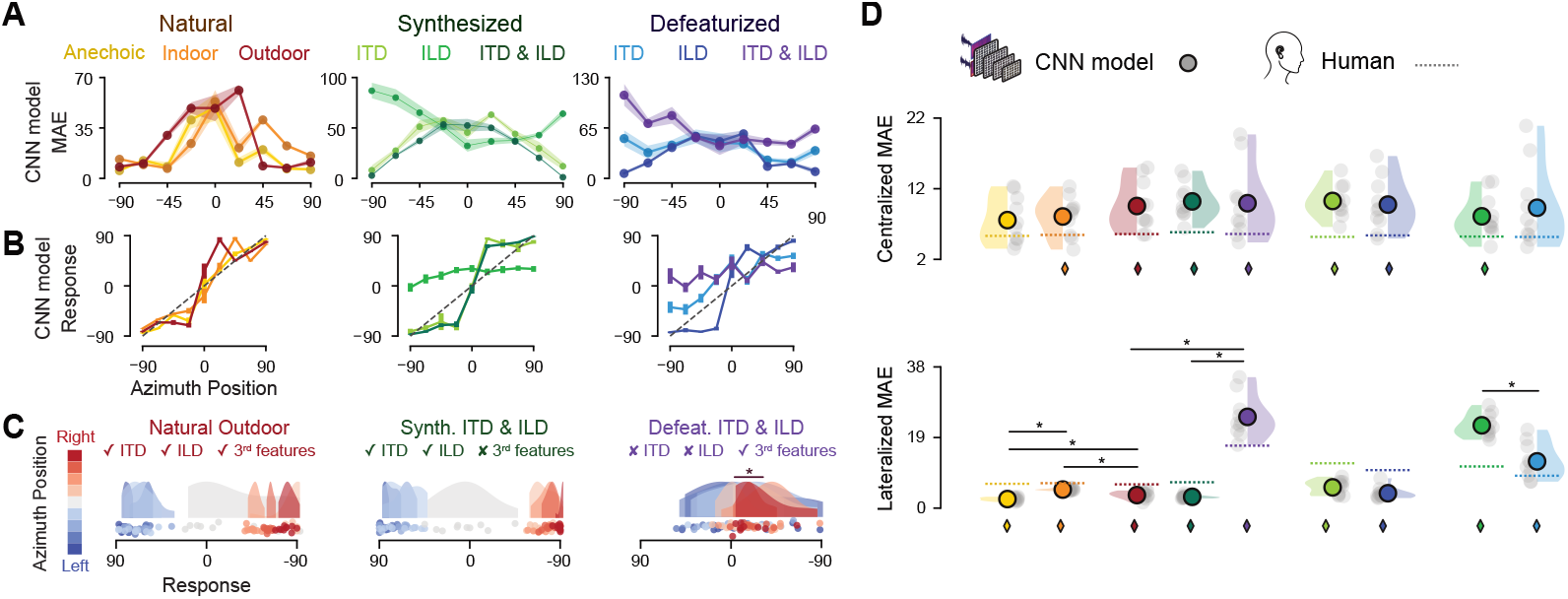
Deep neural networks localization performance and comparison with human behavior. **a**. Lineplots depicting the mean absolute error (MAE) in degrees between the true azimuth and the model predictions as a function of the azimuth position across stimulus types. Shaded areas represent SEM. **b**. Lineplots showing the transfer function of the average CNN model performance mapping the true azimuth position (x-axis) to the responded one (y-axis). Dotted black lines indicate ideal pattern, while horizontal bars indicate SEM across models. **c**. Model prediction distributions for the natural outdoor, synthesis ITD-ILD and defeaturize ITD-ILD stimuli. Each dot represents the average response of each model to a specific location, color-coded by the azimuth position as expressed by the colorbar. Asterisks represent statistical significance. **d**. Raincloud plots showing the centralized (top) and lateralized (bottom) MAE score in degrees of CNN model predictions. Dotted lines represent the average human performance. Asterisks represent statistical significance across stimulus types when comparing model performance, while diamonds indicate statistical significance between human and model performance.

As for humans, we examined the distribution of model responses (Fig. 5C). Models showed an overall response pattern well aligned with the actual azimuth locations, nearly across all stimulus types except synthesized ILD (Fig. S5A). Crucially, the response pattern in the defeaturized ITD-ILD stimuli appeared severely compromised, although CNNs consistently localized sounds correctly to the right (P_FDR_ = 0.02, d = 1.13). In sum, these findings suggest that the behavior of humans and CNN models was well aligned for natural environments but not for out-of-distribution stimuli, especially when lacking primary interaural cues.

### Comparing human and deep neural network localization performance

Next, we compared CNN and human responses for the same stimuli by means of error metrics. No significant difference was observed in the centralized MAE across all stimulus type combinations (Fig. 5D, all P_FDR_ > 0.05), while the lateralized MAE for defeaturized ITD-ILD stimuli was significantly higher than natural outdoor and synthesis ITD-ILD stimuli, mimicking the result pattern found in humans. However, in contrast to human results, no difference was observed between natural outdoor and synthesized ITD-ILD stimuli (P_FDR_ = 0.189, d = 0.45, 95% CI [-0.3, 1.2]), while a significant difference was observed among all pairwise comparisons between natural environments (all P_FDR_ < 0.031, all d > 0.86), with the anechoic environment having the least lateralized MAE. Crucially, we found partial evidence that CNNs used third spatial features when comparing synthesis ITD with defeaturize ILD (P_FDR_ = 0.067, d = 0.69, 95% CI [-0.11, 1.48]) and synthesis ILD with defeaturize ITD (P_FDR_ < 0.0001, d = 1.71, 95% CI [0.59, 2.84]). Notably, when averaging the MAE across all position (Fig. S5B) we also found the same pattern of results that humans exhibited (all P_FDR_ < 0.011, all d > 1.10), with the additional findings that natural outdoor was significantly lower than synthesis ITD-ILD (P_FDR_ = 0.04, d = 0.78, 95% CI [-0.04, 1.6]) and defeaturize ILD significantly lower than synthesis ITD (P_FDR_ = 0.024, d = 0.92, 95% CI [0.06, 1.77]).

When directly comparing the human and model performance (Fig. 5D), we found that humans significantly outperformed CNN models across nearly all the stimulus types in the central positions (except anechoic and defeaturize ITD, all P_FDR_ < 0.036, all d > 1.24), while models outperformed humans in lateral positions (all P_FDR_ < 0.0001, all d > 0.47) across all natural environments and whenever the ITD feature was present (synthesis ITD-ILD, synthesis ITD, defeaturize ILD). Crucially, human lateralized MAE was significantly lower than models in synthesis ILD and defeaturize ITD (both P_FDR_ < 0.033, both d > 1.16) and, most remarkably, in the defeaturize ITD-ILD (P_FDR_ = 0.002, d = 2.14, 95% CI [1.63, 2.66]). Notably, the average MAE (Fig. S5B) showed that humans significantly outperformed models in all synthesized stimulus types and defeaturized ITD and ITD-ILD (all P_FDR_ < 0.003, all d > 1.04), while models performed better than humans in the anechoic environment (P_FDR_ < 0.0001, d = 1.58, 95% CI [1.16, 2.0]).

Together, these findings show that humans and deep neural network models were overall well aligned in their sound localization performance, but humans outperformed models specifically for out-of-distribution stimuli.

### Absence of ITD and ILD cues misalign human and model performance

The above results on synthesized and defeaturized stimuli suggest that humans may outperform CNN models in the use of third features beyond primary interaural cues. To directly address this, we took a conservative approach and first finetuned the CNN models (Fig. S6A) on the train set of our recorded spatial sound dataset (Fig. 6A, referred as “Finetuned CNN” while the previous reported model is now referred as “Sim-train CNN”), because the pretrained model could be biased by the training regime (i.e., the simulation-reality gap^38^). Also, the performance of the Finetuned CNN was severely compromised for defeaturized ITD-ILD stimuli even though right localization was above chance (Fig. S6B-C; P_FDR_ = 0.02, d = 1.03).

**Figure 6.**
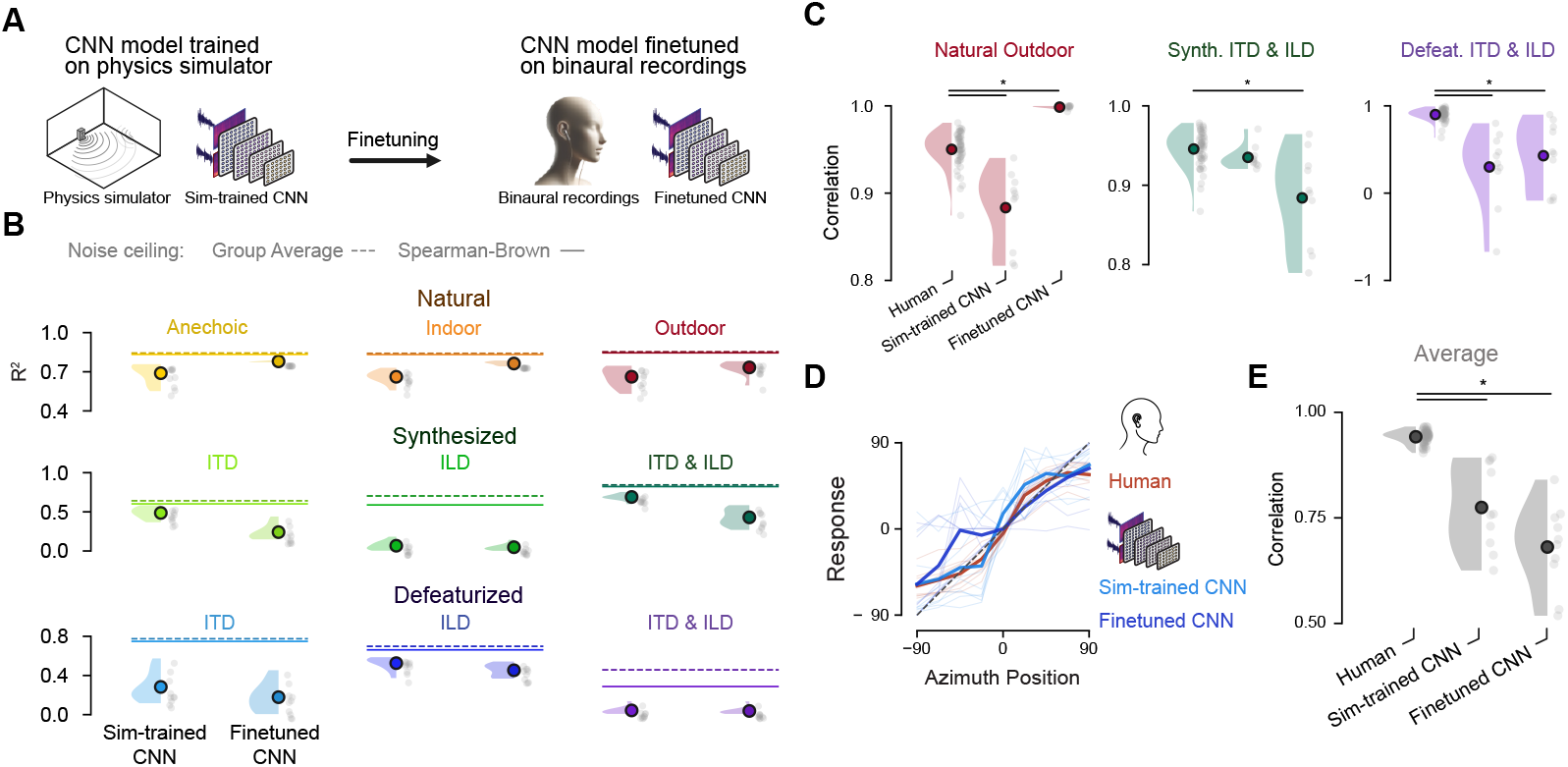
Evaluating human-model alignment and sensory robustness across humans and CNNs. **a**. Schematic illustration of the two version of the CNN models used in this study. The first one, referred as “Sim-trained”, was trained on a virtual physics simulator while the second one, referred as “Finetuned”, was the pretrained model finetuned on the train set of our spatial sound dataset we recorded. **b**. Raincloud plots showing R^2^ scores for linear regression models fitted to predict human responses using as regressor either the Sim-trained or the Finetuned CNN model predictions. Each dot represents the average fit of one model across all human participants, while the circle overlapping the density plot represent the mean across models. Horizontal lines represent noise ceiling estimates, either from the human group average (dotted) or with the Spearman-Brown method (solid). **c**. Raincloud plots showing Pearson correlation values between the ideal performance and human or model responses. Each dot represents individual human participants or CNN model instances, while the circle overlapping the density plot represent the average across humans or models. Asterisks signal statistical significance, generally as P < 0.05. **d**. Lineplot showing the transfer function of the group-average human and model performance mapping the true azimuth position (x-axis) to the responded one (y-axis). Dotted black lines indicate ideal pattern, thin lines the average performance in each stimulus type and the thick lines the average across all stimulus types. **e**. Same raincloud plots as in **c**. showing the Pearson correlation between the ideal performance and the average across all stimulus types.

We next directly quantified how much human behavior variance was explained by model responses (Fig. 6B). We found a strong alignment between models and humans on natural stimuli (noise-corrected R^2^ averaged across environment for Sim-train CNN was 0.80, while for Finetuned CNN was 0.91). Critically, the alignment for synthesized stimuli was strong only when ITD was present (synthesis ITD/ITD-ILD noise-corrected R^2^ for Sim-train = 0.78/0.83, Finetuned = 0.42/0.55), but very low when both ITD and ILD were absent in defeaturized stimuli (noise-corrected R^2^ for Sim-train = 0.11, Finetuned = 0.12). Notably, the noise-corrected R^2^ was significantly lower than 1 (all P_FDR_ < 0.0001, all d > 1.8) in both model sets across all stimulus types. Thus, alignment between human and model behavior specifically broke down for those stimuli for which humans only used third features.

### Robustness of spatial hearing beyond primary interaural cues

To summarize the above findings into a single quantity, we computed the correlation between the average response pattern across azimuth positions and the ideal performance pattern (Fig. 6C). In other words, we quantified to what extent the behavioral transfer function (compare Fig. 2C-5B) approximated the ideal diagonal. For the natural outdoor environment, humans exhibited a remarkable correlation of 0.95 with optimal performance, which was significantly higher than Sim-train CNN (average correlation of 0.88, P_FDR_ = 0.003, d = 2.63, 95% CI [2.04, 3.22]) but significantly lower than the Finetuned CNN model (average correlation of 0.99, P_FDR_ < 0.0001, d = 2.59, 95% CI [2.01, 3.18]). Humans also exhibited a noteworthy correlation of 0.95 in the synthesis ITD-ILD stimuli, which was significantly higher than the Finetuned CNN (correlation of 0.88, P_FDR_ = 0.011, d = 1.96, 95% CI [1.48, 2.44]), but not different from the Sim-train model (correlation of 0.94, P_FDR_ = 0.082, d = 0.49, 95% CI [0.19, 0.79]).

Crucially, humans maintained a high correlation of 0.90 even for defeaturized ITD-ILD stimuli (Fig. 6C), while CNNs considerably dropped in their performance, with a significantly lower correlation of 0.30 and 0.42 as compared to humans for Sim-train (P_FDR_ = 0.004, d = 3.33, 95% CI [2.61, 4.04]) and Finetuned models (P_FDR_ = 0.005, d = 2.9, 95% CI [2.26, 3.54]), respectively. Notably, humans significantly outperformed both model sets on all the remaining non-natural stimulus types except the Finetuned model for defeaturized ILD stimuli and the Sim-train CNN on the natural indoor environment (Fig. S7A, all P_FDR_ < 0.026, all d > 1.46), while the Finetuned CNN exhibited a significantly higher performance compared to humans on the remaining natural environments (both P_FDR_ < 0.0001, both d > 2.23).

Finally, we averaged the performance metric across all stimulus types in order to quantify how robust localization performance was in humans and models across both in-distribution and out-of-distribution scenarios. We found that the average human performance was remarkably close to ideal performance (Fig. 6D), exhibiting a strong correlation of 0.94 across all stimulus types (Fig. 6E). Moreover, humans significantly outperformed CNN models which scored an average correlation of 0.77 (P_FDR_ < 0.0001, d = 4.03, 95% CI [3.18, 4.88]) and 0.68 (P_FDR_ < 0.0001, d = 6.31, 95% CI [5.03, 7.6]) for Sim-train and Finetuned, respectively. Together, these results underscore the robustness of human spatial hearing beyond primary interaural cues and the scarce generalization abilities of current deep network models optimized for sound localization, particularly in out-of-distribution conditions.

### Relevant acoustic features for localization beyond ITD and ILD

What are the spectral characteristics of the third features that enabled in particular humans to localize sounds beyond ITD and ILD? In a first step, we directly contrasted the power and phase of maximally lateralized stimuli presented to both ears (Fig. 7A). This revealed substantial spectral differences in both the power and phase spectra (Fig. 7A). To investigate the contribution of these spectral characteristics to localization performance, we conducted a second behavioral experiment using a two-alternative forced-choice (2AFC) sound localization task (Fig. 7B), in which humans and CNNs were presented with binaural defeaturized ITD-ILD stimuli, along with modified versions that further lacked specific spectral features. Specifically, we matched either the power or phase spectrum across ears (Fig. 7C) or we matched both to remove all interaural cues (leaving only monaural information). This resulted in four different stimulus types tested in the 2AFC task.

**Figure 7.**
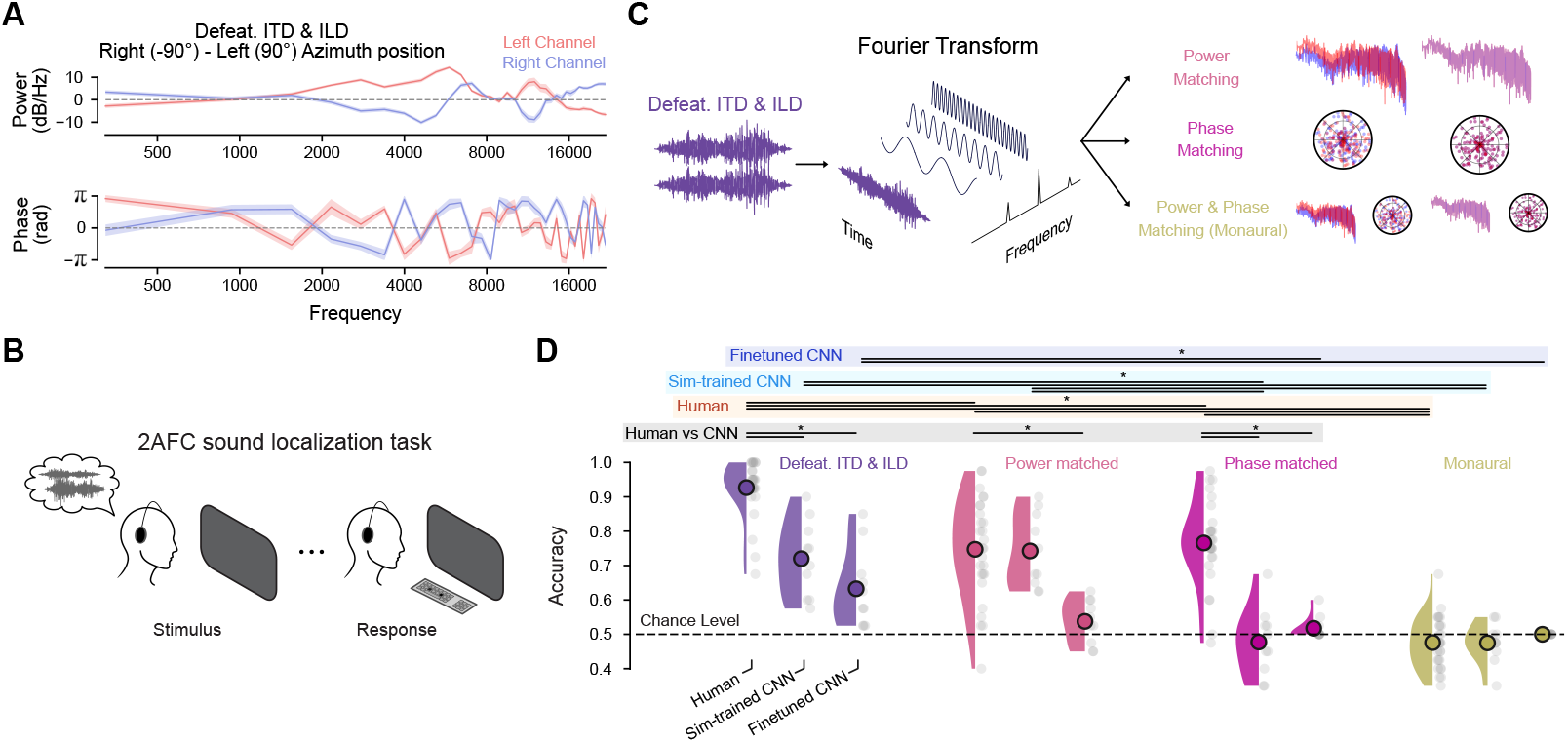
Robustness of spatial hearing in humans and CNN models beyond primary interaural cues. **a**. Lineplots showing the power and phase spectra of defeaturize ITD-ILD stimuli. Each line indicates the difference between the right most and left most azimuth position for each channel. Shaded area represents the standard error of the mean across stimuli. **b**. Schematic illustration of the experimental design adopted for the second behavioral experiment where human participants performed a two alternative forced choice (2AFC) sound localization task. **c**. Graphical illustration of the procedure for generating the stimulus set we used for the 2AFC sound localization task. Starting from the defeaturize ITD-ILD stimuli, we matched the power or phase spectrum (or both) between left and right channels. **d**. Raincloud plots showing the accuracy of humans or CNN models in the 2AFC sound localization task across stimulus types. For power, phase and monaural matched stimuli, the accuracy is assessed from the dominant side (e.g., for stimuli located to the left we considered the accuracy from the generate stimuli with the left channel as reference). Each dot represents individual human participants or CNN model instances, while the circle overlapping the density plot represent the average across humans or models. Horizontal dotted black line indicates chance level performance. Asterisks indicate statistical significance, color-coded by whether the comparisons were tested within human or model performance or between human and models.

We found that humans reported their choices significantly faster for defeaturized ITD-ILD stimuli compared to all the other stimulus types (Fig. S7B, all P_FDR_ < 0.006, all d > 0.59), while monaural stimuli were reported significantly slower compared to power and phase matched stimuli (both P_FDR_ < 0.002, both d > 0.69). Remarkably, human localization accuracy for defeaturized ITD-ILD stimuli was highly significant at 93% (Fig. 7D; P_FDR_ < 0.0001, d = 5.13, 95% CI [3.64, 6.62]), which replicated our results from the main experiment. Moreover, humans performed also significantly above chance for power and phase matched stimuli (both P_FDR_ < 0.0001, both d > 1.68) but not for monaural stimuli (P_FDR_ = 0.104, d = 0.32, 95% CI [-0.08, 0.73]). Humans were significantly more accurate for defeaturized ITD-ILD stimuli than for all its modified versions (all P_FDR_ < 0.0001, all d > 1.45), suggesting that spectral features were indeed exploited for localization. Moreover, accuracy was significantly higher for power and phase matched stimuli than for monaural ones (both P_FDR_ < 0.0001, both d > 1.53), but there was no significant difference between them (P_FDR_ = 0.409, d = 0.16, 95% CI [-0.24, 0.56]; see Fig. S7C for results averaging across both ears). Together, these results suggest that humans consider both interaural power and phase differences for spatial hearing.

Also CNN performance was significantly above chance for defeaturized ITD-ILD stimuli (Fig. 7D) for both models sets (both P_FDR_ < 0.02, both d > 1.15). However, both model sets performed at the chance level for phase-matched and monaural stimuli (all P_FDR_ > 0.176, all d < 0.52). For power-matched binaural sounds only the Sim-train CNN was significantly above chance (P_FDR_ < 0.0001, d = 2.27, 95% CI [0.92, 3.63]). Both model sets performed significantly better for defeaturized ITD-ILD compared to phase-matched (both P_FDR_ < 0.015, both d > 1.29) and monaural (both P_FDR_ < 0.015, both d > 1.15) stimuli but not compared to power-matched (both P_FDR_ > 0.108, both d < 0.7) stimuli. Additionally, the Sim-train CNN’s accuracy was significantly higher for power-matched stimuli compared to both phase-matched and monaural stimuli (both P_FDR_ < 0.002, both d > 1.62), while this was not the case for the Finetuned model (both P_FDR_ > 0.144, both d < 0.59; see Fig. S7C for averages across both ears). These results confirm our previous finding that CNNs are moderately sensitive to third features beyond ITD and ILD and suggest CNNs rely more on interaural phase differences than on power differences or monaural cues.

Finaly, we directly compared the performance of humans and deep neural networks. Humans outperformed models in localizing defeaturized ITD-ILD stimuli (Fig. 7D, both P_FDR_ < 0.0001, both d > 2.27) with a remarkable difference of at least 20% of accuracy with respect to the best model set. Humans also outperformed CNNs on phase-matched stimuli (both P_FDR_ < 0.0001, both d > 2.29), while on power-matched stimuli humans performed significantly better than the Finetuned models (P_FDR_ < 0.0001, d = 1.6, 95% CI [1.01, 2.2]) but not the Sim-train ones (P_FDR_ = 0.969, d = 0.03, 95% CI [-0.36, 0.43]). No difference was observed between humans and models in the accuracy to localize monaural stimuli (both P_FDR_ > 0.104, both d < 0.38; see Fig. S7C for averages across both ears). In sum, these results demonstrate the flexibility of the human auditory system in exploiting relevant features for sound localization compared to deep neural networks.

## Discussion

Our results offer key insights into the sensory processing underlying human spatial hearing and reveal how deep neural networks optimized for sound localization align or diverge from human behavior in both naturalistic and out-of-distribution conditions.

As expected, human lateralization performance was highest for lateral and center locations and lowest for intermediate locations. This baseline performance validated our experimental design and ensured that experimental manipulations applied to natural sounds are rooted in ecologically valid auditory processing. An interesting finding on the natural stimuli was that the combinations of ITD and ILD better predicted human responses than each cue alone. This suggests that humans, when confronted with real-world auditory scenes, make use of spatial auditory features in a more complex way than previously hypothesized by, for instance, the duplex theory^9,39,40^. Specifically, even though the human auditory system may have a frequency-dependent preference for interaural cue reliance, in natural settings it seems to use as much information as it can extract from the sensory data. Notably, our results showed that the combination of interaural cues explained a substantial portion of the variance in human responses. This finding, combined with the findings on the artificial stimuli synthesized with both ITD and ILD showing a remarkable replication of the response pattern on the natural stimuli, confirm previous research on the importance of these primary interaural cues for human spatial hearing^4–6^.

Importantly, our experimental manipulations allowed us to investigate the contribution of additional auditory features to localization performance. For natural stimuli, the residual variance left after predicting human responses using only primary interaural cues indicated that other, non-primary interaural cues (i.e., third features) also play a role. Directly contrasting artificial stimuli, which differed only by third features, showed that these third features indeed enhance localization performance. To our knowledge, these findings uncover a previously unrecognized role of non-primary interaural cues in human spatial hearing.

Previous reports about the negligible effect of third features on human localization performance^9^, may be explained by the experimental design adopted in previous studies where ITD and ILD were always included among stimulus’s features. In contrast, we asked what happens to human localization performance when both primary interaural cues are absent. We found that localization error increased substantially in particular for lateral positions, confirming how heavily humans rely on primary interaural cues. Nevertheless, this acoustic manipulation did not reduce localization performance to chance level. Instead, the response pattern was compressed (e.g., a 90° source was perceived around 45°) but not completely disrupted by the absence of primary interaural cues. This outcome is in contrast to the longstanding emphasis in the literature on the critical importance of ITD and ILD for sound localization^1,2,9^. Thus, we propose that human spatial hearing is inherently more robust than classical theories centered exclusively on ITD and ILD predict. We speculate that, despite the rarity of complete ITD/ILD loss in everyday settings, natural environments may degrade these cues to a considerable degree, necessitating compensatory mechanisms. Notably, our results show that even in the absence of head movements that typically aid in disambiguating spatial cues^41^, listeners maintain a compressed yet coherent spatial percept in absence of primary interaural cues. Furthermore, these results also underscore the power of stimulus generation paradigms to investigate the representational capacity of sensory systems ^23,42^.

Deep neural networks optimized for sound localization replicated many of the findings on human data, such as the error pattern across azimuth positions in natural environments and the restoration of the natural response pattern by including ITD and ILD in synthesized stimuli. Notably, deep neural networks also showed sensitivity to third features. These findings confirm and extend previous results on the alignment between these models and humans on diverse spatial hearing phenomena^29^. However, models exhibited a strong bias for th ITD cue, resulting in a misalignment with human behavior whenever this feature was absent. More importantly, even after fine-tuning on our naturalistic dataset, these models did not match the human robustness to missing primary interaural cues. Interestingly, models increased their reliance on third features when ITD was absent, suggesting that, similarly to humans, they compensated by exploiting alternative acoustic information whenever the preferred cue was unavailable. Nonetheless, these compensatory mechanisms were far less flexible than those of human listeners, resulting in substantially diminished robustness when primary interaural cues were degraded. Future studies may investigate whether spatial hearing alignment between humans and deep neural networks could be improved by the adoption of more ecological training schemes such as self-supervised predictive learning^43–49^.

We argue that the discrepancies between humans and deep neural networks provide profound insights into the computational mechanisms underlying spatial hearing and, more broadly, human sensory processing. In particular, the lack of robustness observed in these models relative to human listeners likely stems from the models’ well-documented tendency to prioritize simple features^50^. This is referred to as the “simplicity bias”^51,52^, an inductive bias inherently present in the behavior of models due to the optimization algorithm used during training^53– 55^. Although this “shortcut” strategy can be beneficial, helping to avoid overfitting despite overparameterization^50,54,56–58^, it also leads to suboptimal performance on out-of-distribution samples^59,60^, as seen here when models encountered stimuli devoid of ITD and ILD. Once primary interaural cues reliably solve the localization task under training conditions, additional cues that might be crucial in more unusual or degraded scenarios are neglected. By contrast, humans appear to maintain a broader search for relevant cues, even when some are consistently reliable, suggesting a cognitive architecture geared toward preserving functionality under a wider range of acoustic conditions. We speculate that evolutionary pressures^61^ may have favored sensory systems with a more distributed reliance on multiple features, as it confers robustness in unusual environments, even if it occasionally sacrifices peak performance under ordinary circumstances. Indeed, although deep neural networks matched or exceeded human performance in typical natural environments, they fell short whenever the most informative cues were absent. From an evolutionary standpoint, this kind of robust flexibility may be more valuable for survival than narrow optimization for the most common acoustic scenarios.

Finally, we sought to identify the third features by further stripping away spectral interaural cues^62^ from stimuli already devoid of ITD and ILD. Interestingly, even when some spectral interaural cues were removed, humans maintained a remarkable spatial accuracy, surpassing the performance of deep neural networks and reinforcing the idea that human auditory processing can flexibly exploit residual cues. Indeed, participants’ performance remained relatively unchanged whether the stimuli were missing power- or phase-based cues, suggesting that humans can leverage whatever information remains available, a hallmark of their robust sensory strategy. By contrast, some models demonstrated a specific preference for phase information, pointing toward a more temporal cue centric localization strategy, consistent with their strong reliance on ITD. In both humans and models, purely monaural cues were largely unhelpful, although we note that in our stimuli, each ear received identical monaural information, making it difficult to ascertain whether such cues might be more effectively used in other contexts. Further investigation is needed to clarify whether monaural features can, in some circumstances, serve as a fallback mechanism when primary and spectral interaural cues are absent.

Our findings have several implications for engineering and clinical applications in spatial audiology. First, our results highlight opportunities to refine artificial systems, particularly those designed for spatial audio processing^63^ or multimodal models^64–66^, by embedding mechanisms that integrate a wider variety of acoustic information and prioritize robustness to sensory perturbations over efficiency for deploying them in real-world scenarios. Such improvements could benefit robotics^67^, virtual/augmented reality^68,69^, and any domain requiring reliable sound localization in a diverse range of acoustic conditions. Second, these insights are relevant for clinical applications, such as the optimization of cochlear implants^70–74^ and other auditory prosthetics. By incorporating similar computational strategies for extracting spatial cues like the ones uncovered here, these devices may ultimately enhance the user’s spatial awareness and speech intelligibility in noisy or unusual acoustic environments.

In conclusion, our findings provide evidence that human spatial hearing remains remarkably robust even when primary interaural cues are absent. Moreover, the stark contrast with deep neural networks reveals that humans employ a fundamentally different computational strategy to process sensory data for spatial hearing. Our findings deepen our understanding of human auditory processing and point to promising avenues for enhancing artificial systems and optimizing clinical interventions such as cochlear implants.

## Methods

### Binaural sound data recording

We collected a dataset of spatial sounds to finetune DNNs and test humans and DNNs on sound localization. We recorded these sounds using binaural recording microphones placed inside the ears of a human. This allowed us to faithfully record all the distortions from the human body, including the head and the pinna, that affect the sound waveform up to when it enters the ear. The sounds were played from a speaker located on 9 different positions on the azimuth plane, from 90° (left) to −90° (right) in steps of 22.5° (with the 0° angle positioned in front of the human listener), at 0° elevation with respect to the human listener and the binaural recording system, with a distance of approximately 2 meters separating the speaker from the recording system.

The recordings were performed on 3 different acoustic environments, one anechoic room, one large lecture hall (indoor) and in a garden (outdoor). In each acoustic environment, the human listener wearing the binaural recording system was seated on a chair or a bench, maintaining a forward-facing gaze with minimal head movement. As source sounds, we selected the 50 monaural sounds that were rated as most frequently heard in everyday life by humans from the Natural Sound Stimulus Set dataset^32^. The stimulus duration was 2 seconds and the frequency sample was 44100 Hz for all the selected sounds. All stimuli, for each azimuth position, acoustic environment and stimulus identity, were repeated only once. After the data collection, data were split in one set for training and finetuning the DNNs and one for testing both humans and DNNs. Specifically, the test set was designed to maximize exhaustivity in evaluating the sound localization performance, both in terms of stimulus variability and the contribution of the primary interaural cues.

Thus, we used two criteria to select 10 stimuli out of the total 50 ones that would have been part of the test set. The first one was based on the alignment of the ITD and ILD features to the azimuth position, measured as the Pearson’s correlation between the azimuth angle at which a specific sound was played and the computed ITD or ILD. We used a cutoff of 0.9 in both features as an inclusion criterion. In other words, a sound could be selected for the test set if was correlated at least 0.9 with the correct azimuth angle in both primary interaural cues. We computed ITD as the cross-correlation between the left and right channel. We pad the signal by adding 100 ms of actual recording prior the onset of the stimulus and computed the cross-correlation from −100 ms up to 100 ms after stimulus onset, as follow:

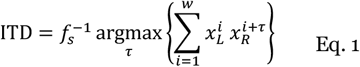

where *x*_*L*_and *x*_*R*_ are the left and right channels, separately normalized to ensure consistent amplitude scaling, *τ* is the time lag (in samples), *f*_*s*_ is the frequency sample and *w* are the number of samples in the selected window above (8820). Negative ITD values indicated that the right channel led the left channel, and vice versa. The ILD feature was computed as the decibel normalized ratio between the root mean square (RMS) of the left and right channel, as follows:

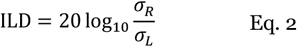

where *σ*_*L*_ and *σ*_*R*_ are the left and right channels’ RMS. The second criterion was based on the “semantic” features of the sounds. We selected the most diverse stimuli that satisfied the first criterion in order to cover the range of sounds that humans usually listen as much as possible. We used the YAMNet^33,34^ model, an audio event classifier deep network model with the MobileNet v1 architecture, to extract the semantic features from the sounds as the embedding vectors from the average-pooled output that feeds into the final classifier layer. Then, we computed a distance matrix by calculating the Euclidean distance between every pair of embedding vectors for each sound. We also plot this matrix using multidimensional scaling (MDS). Note that here we used the original monaural sounds and not the spatial recorded ones, since we were not interested in spatial features. Next, we averaged these distances for each sound and filtered out any that failed to meet our first criterion. Finally, from the remaining set of sounds, we selected 10 sounds whose average distances were evenly spaced across the remaining distance range.

### Human participants

We recruited 52 human participants (32 female, mean age 26.04 years, SD = 4.87) for the main behavioral experiment and 27 participants (10 female, mean age 30.66 years, SD = 4.18) for the second experiment. All participants provided informed consent and had normal hearing abilities. The experiment was realized in accordance with the Helsinki Declaration and the local ethics committee of the Medical Faculty of the University of Tübingen approved the study (No. 014/2022BO1).

### Stimulus generation

To investigate the role of interaural time difference (ITD) and interaural level difference (ILD) in spatial hearing, we created artificial stimuli to complement the naturally recorded sounds in our spatial sound “test” dataset. Specifically, we generated two additional sets of stimuli, referred to as “synthesis” and “defeaturize”. The synthesized stimuli were produced by adding either the ITD, ILD, or both to monaural source sounds. The specific ITD and ILD values were drawn from actual recordings of natural sounds, particularly those captured in the outdoor environment. For the ITD synthesized stimuli, we created a binaural sound by shifting in time the original monaural sound in one channel with respect to the other. For the ILD synthesized stimuli, we generated a binaural sound by normalizing the original monaural by its standard deviation and then multiply the left and right channel with the respective RMS values of the recorded sounds’ channels. This allowed us a direct comparison with the natural stimuli, at least with the outdoor natural stimuli, since they possessed the same primary interaural features but the synthesized stimuli lacked all the other third features. By “third features”, we refer as any acoustic property present in binaural recordings that is neither ITD nor ILD, including monaural cues (e.g., pinna effects) and interaural spectral differences.

In contrast, the defeaturized stimuli were created by “removing” the ITD, ILD, or both from the naturally recorded outdoor sounds. For the ITD defeaturized stimuli, we generated a binaural sound starting from the recorded natural outdoor sounds, by shifting in time back or forward a channel with respect to the other according to the actual ITD in that recorded sound. In other words, this procedure generated a binaural sound that has no time delays between channels. For the ILD-defeaturized stimuli, we divided each channel of the recorded natural outdoor sounds by their own RMS, in order to obtain a binaural sound with the same loudness across the left and right channels. In sum, the natural sounds contained ITD, ILD, and third features, the synthesis stimuli lacked the third features, and the defeaturize stimuli lacked ITD and ILD.

Finally, for the second experiment, we also generated variants of the ITD-ILD defeaturize sounds to further investigate the impact of the remaining acoustic features on the localization performance. We generated power and phase matched stimuli by computing, for each ITD-ILD defeaturize sound, the discrete Fourier transform on the left and right channel separately, converting them into the frequency domain and obtaining their respective magnitude and phase spectra. Then, we designated each channel as the reference channel and replaced the non-reference channel’s phase, power, or both, with those from the reference channel. Specifically, if only phase was matched, we preserved the original magnitudes while substituting the phase values. If only power was matched, we retained the original phases while substituting magnitudes. For complete matching, both the phase and power of the non-reference channel were replaced with those of the reference, effectively copying one channel from the other. After this procedure, we transformed both channels back to the time domain using the inverse Fourier transform. This process effectively allowed us to control or eliminate any interaural differences in phase and or power.

### Behavioral tasks

For the main experiment, we probed human participants with these natural, synthesize and defeaturize stimuli to assess their sound localization performance. The experiment was implemented in Python using the PsychoPy (version 2022.1.4) library^75^. Participants were sitting in front of a computer screen and stimuli were delivered via headphones. They were instructed to perform a sound localization task, in which the presented sounds could come from any location (even though only nine specific locations were presented) in the frontal side of the azimuth plane at 0° elevation. Audio volume was regulated based on individual preference, with the objective to clearly hear a test sound which was not present in the experimental stimuli. The stimuli were all the natural sounds in the test dataset described above, with the additional synthesis and defeaturize counterparts sounds. In total, there were 810 sounds, since there were 10 different sounds and each stimulus type had 3 variants (natural stimuli had different environments, synthesis and defeaturize different contribution of the primary interaural cues) with 9 azimuth locations. All the stimuli were presented once and in random order. Each trial started with a visual cue (a speaker icon) signaling the incoming of the acoustic stimulus, which lasted 2 seconds. Then, participants reported their response on a continuous scale from −90 to 90 azimuth degrees, using the mouse to move a bar on the screen indicating the response angle on the azimuth plane relative to the head of the listener. After they provided the response, they were asked to confirm their choice. In the negative case, they repeat the response. Then, they were asked to provide a confidence score for their response on a slider with a scale ranging from 1 to 5, with the first, third and last score textually labelled as “No idea”, “I guess so” and “Absolutely”, respectively. Finally, after 0.5 seconds of inter trial interval (ITI), a new trial started. Notably, no feedback about the correctness of the response was given. Every 80 trials, the task was interrupted to let participant rest if they needed and then restart the experiment. Prior to the experiment, we administered a frequency discrimination test to confirm that participants could perceive the typical human hearing range. The test consisted of 18 trials presented in random order. In half of them, we presented 9 sine waves linearly ranging from 200 to 10000 Hz, each lasting 1 second, and in the remaining trials, we presented no sound. Participants were asked to indicate whether they heard a sound. All participants passed the test successfully. Importantly, before the start of the sound localization task, participant familiarized with the task in a practice session. We presented 3 sounds from the train set of the recorded spatial sound dataset, which were not included in the main task. The sounds were selected from the natural anechoic environment across all the available azimuth positions. No stimuli were presented belonging to the synthesis and defeaturize set. Thus, a total of 27 stimuli were presented. Crucially, we provided feedback after every trial to indicate whether the participant’s response was correct. This feedback was necessary because a prior pilot study revealed that participants often chose extreme lateral positions even though they were instructed that sound could originate from any direction. By offering a dedicated practice session with feedback, we aimed to reduce this bias and emphasize that the sound could come from intermediate azimuth locations as well.

For the second behavioral experiment, we administered a two alternative forced choice (2AFC) sound localization task. Participants completed the task on a laptop and stimuli were delivered through headphones. We instructed them to report if the sound they heard at each trial were coming either from the left (90°) or the right (−90°) in the azimuth plane, by using two keys to press on the keyboard with their left and right finger. The stimuli were all the 10 sounds from the ITD-ILD defeaturize set, with additional stimuli generated from these as described above in the stimulus generation section. The stimulus set was comprised of ITD-ILD defeaturize (the same used in the main experiment), power-matched, phase-matched and monaural sounds. Thus, there were in total 7 different stimuli “classes”, since the latter three were having the reference channel either from the left or from the right. All the stimuli were presented twice and in random order, resulting in a total number of presented stimuli equal to 280. Each trial started with a visual cue (a text message) signaling the incoming of the acoustic stimulus, which lasted 2 seconds. Then, participants reported their 2AFC response and after 0.5 seconds of ITI, a new trial started. No feedback about the correctness of the response was provided.

### Model architecture, training and testing

A key goal of this study was to compare the human localization performance with modern deep neural network models optimized for sound localization. To this end, we employed convolutional neural networks (CNN) pretrained^29^ in a virtual acoustic world simulator^76,77^ and tested their performance on the same stimuli presented to humans. These models can be viewed as ideal observer models, trained to discover an optimal solution for a real-world task like sound localization, which lacks a simple or analytically tractable solution. Although these pretrained CNNs were not designed to have exact biological correspondence with neural systems in terms of internal representations^29^, they still possess a level of biological plausibility that is either comparable or higher than classical computational models of sound localization ^78,79^ or current engineering systems designed to perform sound localization^80,81^. Importantly, these models have demonstrated remarkable performance at localizing sound under realistic conditions and exhibit many of the same features observed in human spatial hearing, such as sensitivity to monaural spectral cues, ITD and ILD, integration across frequencies and limitations in localizing multiple concurrent sources^29^.

For a full characterization of the model architecture and training details, we refer readers to the original paper^29^. Here, we provide a self-contained description of the key aspects of the model architecture and training process relevant to this study. The CNN model architecture consists of a series of hierarchically organized convolutional block layers, followed by fully-connected layers for the final prediction of the sound location. The input of the model was a cochleagram of the binaural sound waveform. This cochleagram is a biologically plausible time-frequency representation of the human cochlea output simulating the instantaneous mean firing rates in the auditory nerve. The cochlea model^35^ comprised a bank of 39 bandpass filters, with center frequencies ranging from 45 to 16975 Hz, that captured the frequency selectivity of the human ear. The waveforms for the left and right channels were first upsampled to 48 kHz, then separately passed through the filter bank. Filtering was performed in the frequency domain, yielding subbands processed with a power function (0.3 exponent) to simulate outer hair cell compression. These were half-wave rectified to mimic auditory nerve firing rates, low-pass filtered (4 kHz cutoff, Kaiser-windowed sinc with 4,097 taps), and downsampled to 8 kHz. This preserved information across all audible frequencies with fidelity limits consistent with those in the human ear, since the filtering and downsampling were applied to rectified outputs. Thus, the resulting cochleagram had a dimensionality of 39 frequency channels × 8,000 samples at 8 kHz × 2 ears, since the duration of the inputted sound was fixed at 1 second. Next, a convolutional block layer^10^ was typically defined as a sequence of operations comprising a 2D convolution, 2D max-pooling, rectified linear units (ReLU)^82^ activation function and batch normalization^83^. Finally, after the convolutional blocks, there were always two fully-connected layers. The first one was consisted of 512 units, followed by a ReLU activation function, batch normalization and a dropout layer^84^ with 0.5 probability. The last one was the output layer with 504 units and a softmax activation function, corresponding to different spatial locations equally spaced 5° in azimuth and 10° in elevation.

These models were trained to perform sound localization in a virtual environment, casted as a classification problem with the cross-entropy loss function and stochastic gradient descent (SGD) to optimize model weights. The training samples were derived from 385 natural sound recordings^32^ rendered spatially in a simulated room. Using a modified source-image method^76,77^, the room simulator modeled wall reflections, with each reflection filtered by direction-specific binaural head-related impulse response (HRIRs). Five rooms with varying dimensions and wall materials introduced environmental variability, and spatialized noise was added to replicate real-world auditory conditions. Model hyperparameters, such as the kernel width and height of the convolutional or pooling layer, were optimized via architecture search, which involved training a set of 1500 randomly sampled candidate architectures for 15000 steps with a total of 240000 samples. The 10 best-performing networks on a validation dataset were selected for further refinement. Their parameters were reinitialized, and each network was trained for 100000 steps with 1.6 million samples.

In our study, we used these 10 optimized CNN models, treating them analogously to individual participants in a behavioral experiment, as we did with our human participants. Additionally, we finetuned each of these models on the train set of our spatial sound dataset we collected. This train set consisted of 40 different binaural sounds from 9 azimuth positions and 3 environments (see the above section “Binaural sound data recording” for details), resulting in a total of 1080 samples. Thus, we retrain these CNNs starting from their trained model parameters for 50 epochs and using a batch size of 32 and as optimizer Adam^85^ with a learning rate of 0.001, β^1^ as 0.9 and β^2^ as 0.999. On each epoch, we applied data augmentation strategies by randomly slicing the binaural sound samples in the temporal dimension and randomly leveling the audio volume. The original duration of the samples was 2.2 seconds (2-second waveform duration with a 0.1 second padding), from which we selected 1 second since the CNN expected this temporal dimensionality. Moreover, the volume was randomly adjusted from 0.75 to 1.25 of the original audio volume of the sound samples. This data augmentation procedure helped the model to regularize the training process by preventing it to focus on unreliable features given by the onset of the audio input or its overall audio intensity. Since we presented these finetuned version of the CNN models in the results section, we referred to the pretrained models as “Sim-train CNN” and to these ones as “Finetuned CNN”.

We tested these CNN models on the same stimuli we used to probe humans’ sound localization performance. Thus, we collected their localization predictions on the natural, synthesis and defeaturize sounds using a sliding window approach. We opted for this since the model had a fixed input dimensionality of 1 second waveform and our experimental design comprised of 2 seconds stimuli. To fairly compare humans and deep network models, we averaged the predictions from the models across time using 3 windows of 1 second with 0.5 seconds of overlap. Importantly, we only collected the predicted azimuth position from the models and ignored the predicted elevation angle, by additionally clipping the azimuth response to −90 to 90 degrees for comparability purposes with the human responses.

Additionally, we also tested the hidden representations of the models by investigating the predictive performance on the azimuth location across the layer depth. Thus, for each model, we extracted the hidden representations, with the same sliding window approach as above, defined as the layer activation after the 2D convolutional operation in each convolutional block. We then reduced the dimensionality of these layer activations by concatenating all the stimuli’s representations across the sample dimension and retained the first 100 dimensions obtained via Principal Component Analysis (PCA). Localization performance was assessed using a Support Vector Regression model with a 5-fold cross-validation scheme, radial basis function as kernel and a regularization parameter of 1 and plotted across layers by interpolating the absolute layer depth index of each model in a normalized layer depth range. For visualization purposes, we also plotted the first two principal components for each stimulus type and convolutional block on an exemplar CNN.

Moreover, we investigated the models’ hidden representations using a “system neuroscience” approach^36,37^, aiming to identify and compare artificial units’ feature selectivity across stimulus types. This approach consisted of contrasting the units’ activity across stimulus types, as usually done in neuroscientific studies by comparing neurons’ spiking activity across experimental conditions. Thus, we used the same layer activity extracted in the previous analysis and performed critical comparisons to elucidate if an artificial unit was sensitive to the ITD, ILD or third features. Each unit activity was compared between stimulus types by averaging their difference across stimuli and dividing by the standard deviation of their difference, essentially z-scoring their comparison. Then, we averaged the number of significant units per layer, defined as those who passed the threshold value of a two-tail z-test with an alpha of 0.05 (circa ±1.645), and plotted it in a normalized layer depth range.

### Data analysis

We started analyzing human data by preprocessing their responses based on their general performance. We averaged the mean absolute error (MAE) between human responses and actual azimuth position across all stimulus types and used as outlier detection criterion the threshold of the median MAE of the whole sample plus the five times the median absolute deviation (MAD). This led us to exclude one participant, which had an average MAE of 65° while the group-average was 22°, resulting in a final human sample of 51 participants. Thus, we plotted the human responses as a function of the azimuth position in a 2D probability density plot and as a transfer function mapping the average responses at each azimuth position, with the ideal performance standing on the diagonal of the plot. Then, we visualized the MAE, accuracy, reaction times and confidence scores as a function of the azimuth position to qualitatively inspect the overall human performance. Next, we correlated the confidence scores with the absolute value of the ITD and ILD cues, to investigate whether the confidence that participants reported was influenced by the magnitude of the primary interaural cues.

Then, we tested the specific contribution of ITD and ILD to the human spatial hearing performance by predicting human responses via linear regression using as regressor either ITD, ILD or their combination. Predictive power of the linear regression model was evaluated by means of the R^2^, adjusted-R^2^ and MAE metric. We also computed two estimations of the noise ceiling to properly compare the R^2^ values independently of the noise level present in the data. One noise ceiling estimation was carried out by averaging the responses of all participants and use them to predict individual participants on each stimulus type. We reported the average of this as the “Group Average” noise ceiling estimate for each appropriate metric. The other noise ceiling estimation procedure was the Spearman-Brown^86,87^, which was computed by randomly half-splitting the ten trials in each stimulus type 100 times and computing the correlation between the two halves with the Spearman-Brown correction. After squaring each value, we reported the average of this procedure and across participants as the “Spearman-Brown” noise ceiling estimate. We reported this noise ceiling only for the R^2^ results and not for the MAE metric since this procedure is based on the correlation values that can be only compared with the coefficient of determination after the squaring operation. Moreover, we derived a noise-corrected version of the R^2^ by dividing it by the average of the two noise ceiling estimates, providing a more realistic upper bound for evaluation than using the absolute value of 1. Similarly, we also fitted the human responses to the artificially generated stimulus types to the natural outdoor responses for each participant to investigate the similarity between the response patterns at these artificial stimuli that specifically manipulated ITD and ILD cues and the response patterns at the natural stimuli.

Next, we computed a metric of central and lateral performance by using a weighted average of the MAE across the azimuth positions to quantitatively investigate the human performance across these stimulus types. Specifically, we defined a scaled exponential function where the weight at each position was computed as *w*_*i*_ = *τ*^|*i*−|^, where *w*_*i*_ represents the weight at position *i, c* is the central position within the defined range (here 5 since there were 9 positions), and *τ* is a scaling factor that controls the rate of decay. Here, we set *τ* to 0.3, meaning weights decreased exponentially as they moved away from the center. Weights were normalized relative to the center position. The center-weighted kernel was directly derived from this function, while the lateral-weighted kernel was constructed by symmetrically mirroring and concatenating weights from the lateral positions, excluding the central region. For completeness, we also computed and reported the average MAE across all positions. Moreover, we computed the distance matrix and its 2D projection using Multidimensional Scaling (MDS) between all stimulus types by computing the Euclidean distance between their MAE pattern across all azimuth positions for each participant and then average the results. Importantly, we also investigated whether the absence of ITD and ILD cues could lead to random responses or if the remaining acoustic spatial features could still enough information for basic localization by computing the average response for each azimuth position and participant and plotted the response distributions as a function of the actual location fro all stimulus types.

*T*hus, we repeated these analyses on the CNN model responses we collected on the same stimuli to compare human and deep neural network behavior. We computed the transfer function and MAE across azimuth positions, as well as quantify the model central and lateral performance by means of the center-weighted and lateral-weighted kernel. Additionally, we used linear regression to predict human responses from Sim-train and Finetuned model responses on all stimulus types, using R^2^ to quantify the predictive performance of the linear regression model and the same noise ceiling estimation procedures as above. Crucially, we also assessed the response distributions of the models with the same procedure described for the human data. Importantly, we defined a measure to summarize the localization performance of humans and models, by computing the Pearson correlation coefficient between the average response pattern across azimuth positions and the ideal performance pattern for each human participant and model instance. Essentially, this allowed us to measure how close was the localization performance to the optimal behavior, by comparing the transfer function we previously computed and plotted to the diagonal line representing the ideal performance. Additionally, we average all the transfer function computed on the stimulus types within each human participant and model instance to obtain a single value, which captured the robustness of the performance to all the sensory perturbations we implemented in our experimental design.

Finally, we analyzed the results from the 2AFC sound localization task. We firstly plot the reaction time probability distributions for each stimulus type and computed the mean reaction time for each human participant. Then, we computed the accuracy of the responses as the percentage of the responses that matched the correct location for each human participant. For the CNN models, binary choices were collected as the responses that exceeded or subceeded the 0° azimuth response and accuracy was defined as the percentage of choices matching the correct location. To put it simply, if the correct location was left and the model responded with an azimuth angle between 5 and 90 degrees (since the model response resolution was 5°) we counted this response as correct and assigned 1 to that response and 0 if it was incorrect. When the model responded exactly at 0° azimuth, we assigned 0.5 to that response effectively counting as chance. Thus, accuracy was computed as the average of these assigned values. Notably, for the power-matched, phase-matched and monaural stimulus types the accuracy values were computed for the dominant side and for both sides. For the dominant side, we considered only the trials in which the spectral features were matched with respect to the reference channels that was on the same side of the actual location. In other words, if the true location at the current trial was left we only considered countable that trial if the right channel was matched to the left one. This was because the opposite trial has the theoretical disadvantage that the relevant channel for sound localization is the most proximal to the sound source. For both side accuracy, we computed the accuracy as an average of both sides.

### Statistical analysis

Statistical analyses were carried out using two-tail t-tests with a significance threshold α set to 0.05. We used paired tests to compare stimulus types within human participants and model instances and independent sample tests to compare human and deep neural network performance. Cohens’d was used as a measure of effect size and reported alongside its 95% confidence interval (CI), with the lower and upper values given by d value plus or minus the standard error of the mean (SEM) multiplied by the critical value of the T distribution corresponding to the selected confidence level. Multiple comparison correction was performed using the False Discovery Rate (FDR) correction^88^, by considering all the possible comparisons were available on each measure of interest.

## Acknowledgements

This study was supported by the European Research Council (ERC; https://erc.europa.eu/, CoG 864491 to M.S.), by the German Research Foundation (DFG; https://www.dfg.de/, projects 276693517 (SFB 1233; to M.S.) and SI 1332/6-1 (SPP 2041; to M.S.)) and by the Horizon Europe programme under the Marie Skłodowska-Curie Actions (MSCA; https://marie-sklodowska-curie-actions.ec.europa.eu/, CherISH Doctoral Network 101120054; to C.B.).

## Author contributions

**A.G**.: conceptualization, software, methodology, investigation, formal analysis, validation, data curation, visualization, writing - original draft, writing - review and editing.

**S.B**.: conceptualization, software, methodology, investigation, formal analysis, data curation, writing - review and editing.

**C.R**.: conceptualization, investigation, formal analysis, writing - review and editing.

**M.S**.: conceptualization, supervision, resources, project administration, funding acquisition, writing - review and editing.

**C.B**.: conceptualization, supervision, resources, project administration, funding acquisition, writing - review and editing.

## Supplementary material

**Figure S1.**
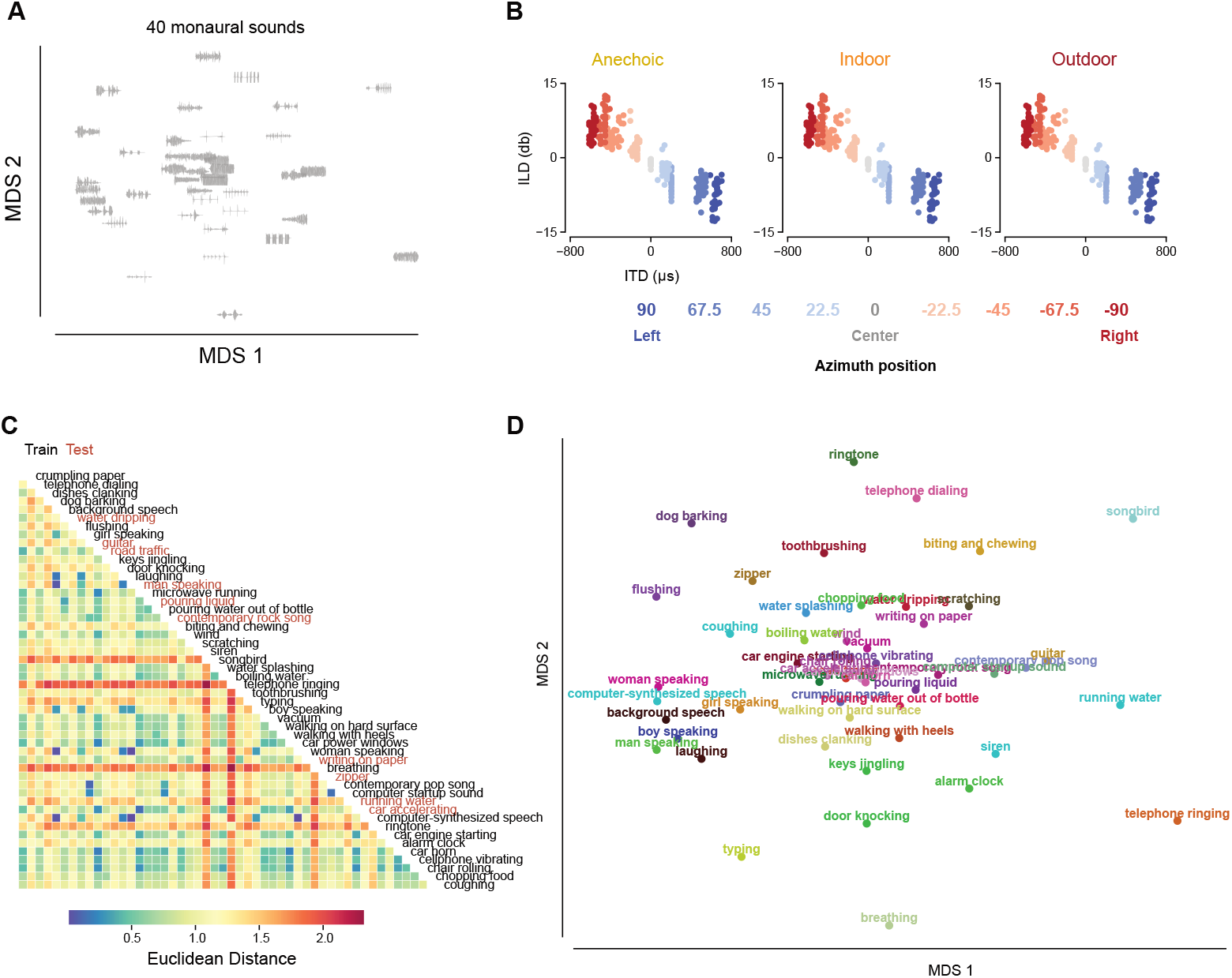
Training set of the spatial sound dataset. **a**. Of the 50 sampled monaural sounds from the Natural Sound Stimulus Set dataset, we used 40 of them to train and finetune deep neural networks (DNN) on sound localization. Here, we projected these sounds on a 2D plane using multidimensional scaling (MDS) based on the distance matrix of the sound waveforms. **b**. Scatterplots showing the ITD and ILD of the resulted sounds in this training set across the 3 environments and color-coded as a function of the azimuth position. **c**. Distance matrix of all the 50 monaural sounds we used to record the binaural sounds in the azimuth plane, using the Euclidean distance on the embeddings extracted from the YAMNet model. This model allows us to visualize the sounds’ distances based on semantic features rather than perceptual ones. The labels are color-coded to distinguish the ones we used to test humans and DNNs (red) to the ones we used for training and finetuning DNNs (black). While in **d**., there is the MDS projection in 2 dimensions of the distance matrix.

**Figure S2.**
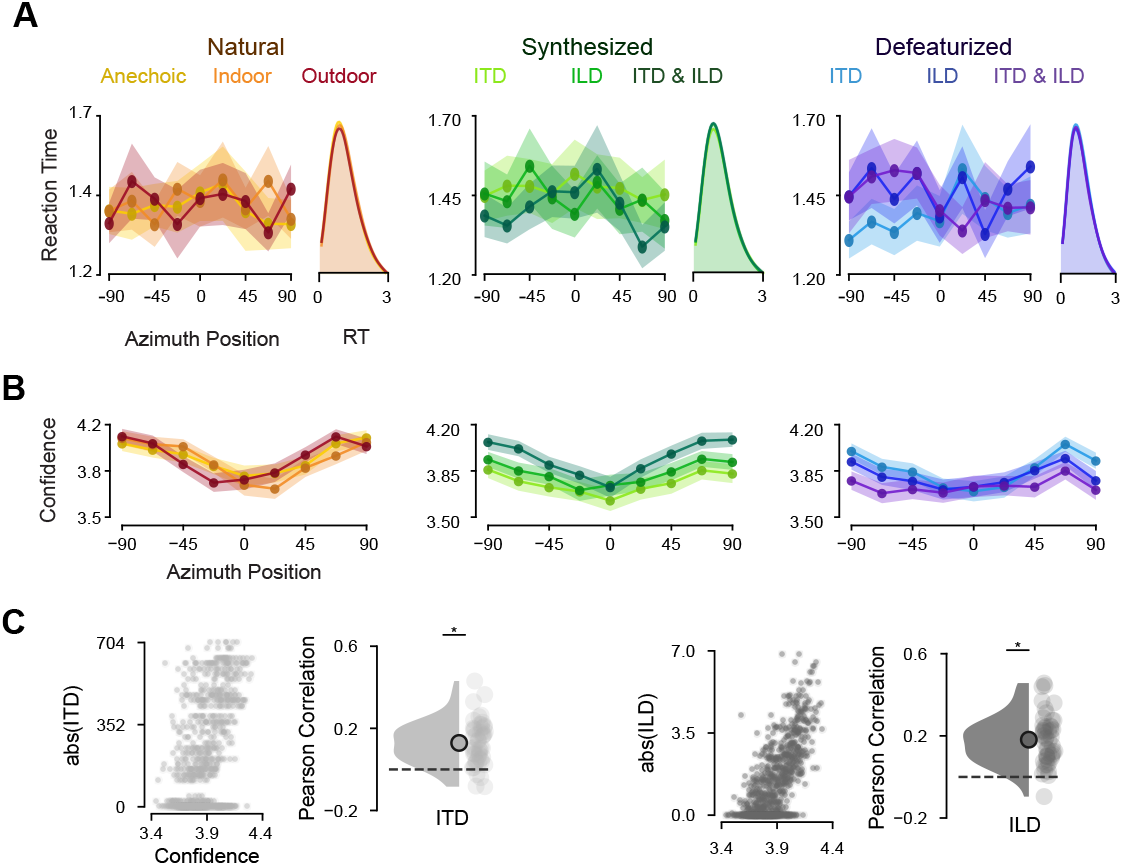
Reaction times and confidence scores from human participants. **a**. Reaction times (RT) of human responses as a function of the true azimuth position across all natural environments and artificial stimulus types. For the three macro types of stimuli (natural, synthesis and defeaturize), on the left there are lineplots showing the average RT (in seconds) for each of the 9 azimuth positions. Shaded areas indicate SEM. On the right, the density plot of the RT distribution (in seconds) across all positions, with all stimulus types largely overlapping. **b**. Confidence scores across the azimuth positions with shaded areas representing SEM. **c**. Correlation between confidence scores and the absolute values of ITD (left) and ILD (right) across all stimuli.

**Figure S3.**
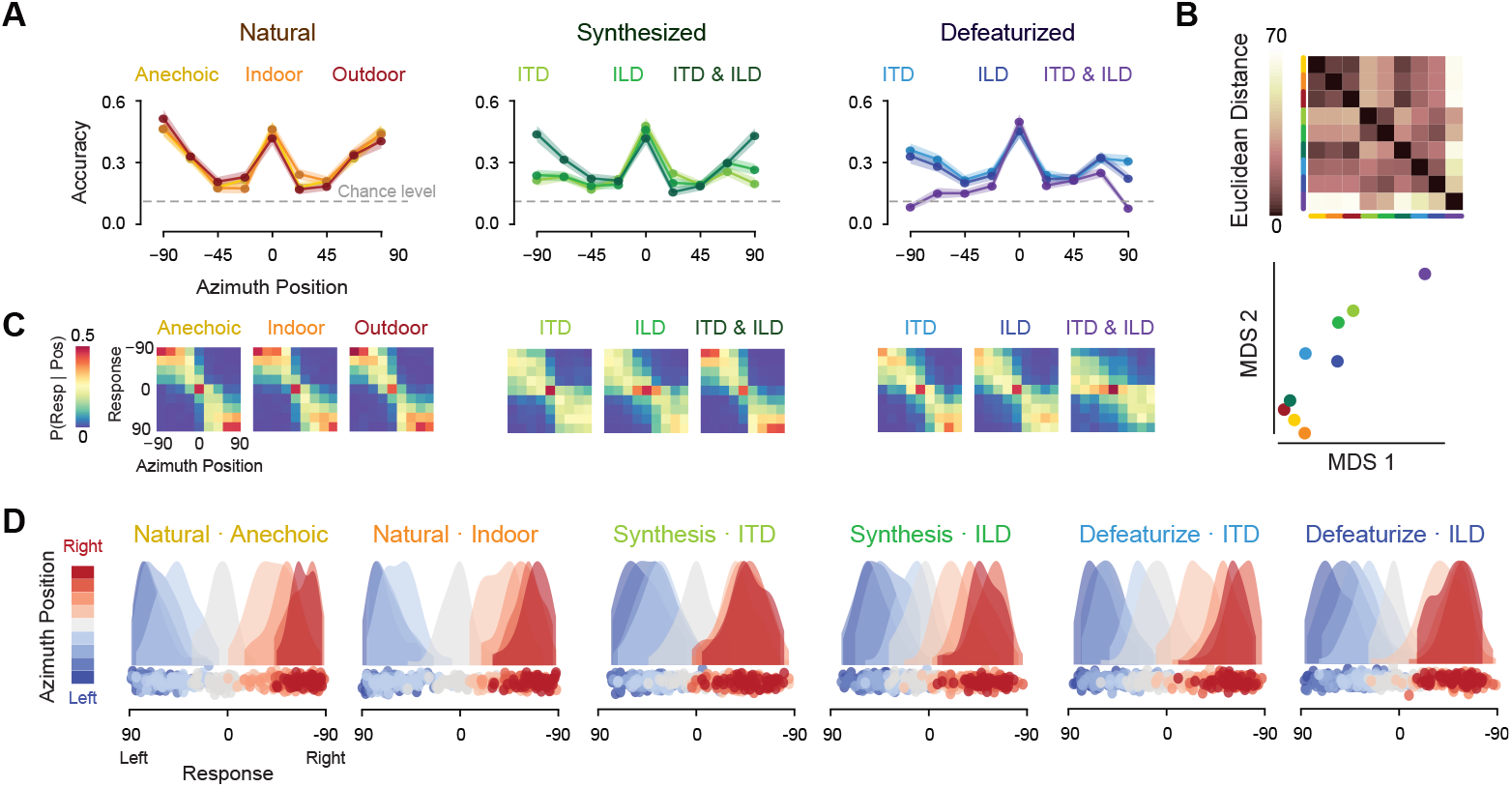
Characterizing human spatial hearing behaviour. **a**. Lineplots depicting the accuracy score between the true azimuth and the human responses as a function of the azimuth position across stimulus types. Dotted gray lines indicate chance level, while shaded areas represent SEM. **b**. On the top, distance matrix between all natural and artificial stimuli. Each cell shows the Euclidean distance between the error patterns across azimuth positions on each pair of conditions. On the bottom, the multidimensional scaling (MDS) plot projecting the distance matrix in 2 dimensions. Dots represent all the conditions. **c**. Distribution of human responses as a function of the true azimuth position on the x-axis and the response position on the y-axis, across all natural environments and artificial stimulus types. **d**. Response distributions for the natural, synthesis and defeaturize stimuli that were not shown in Fig. 2D. Each dot represents the average response of each participant to a specific location, color-coded by the azimuth position as expressed by the colorbar.

**Figure S4.**
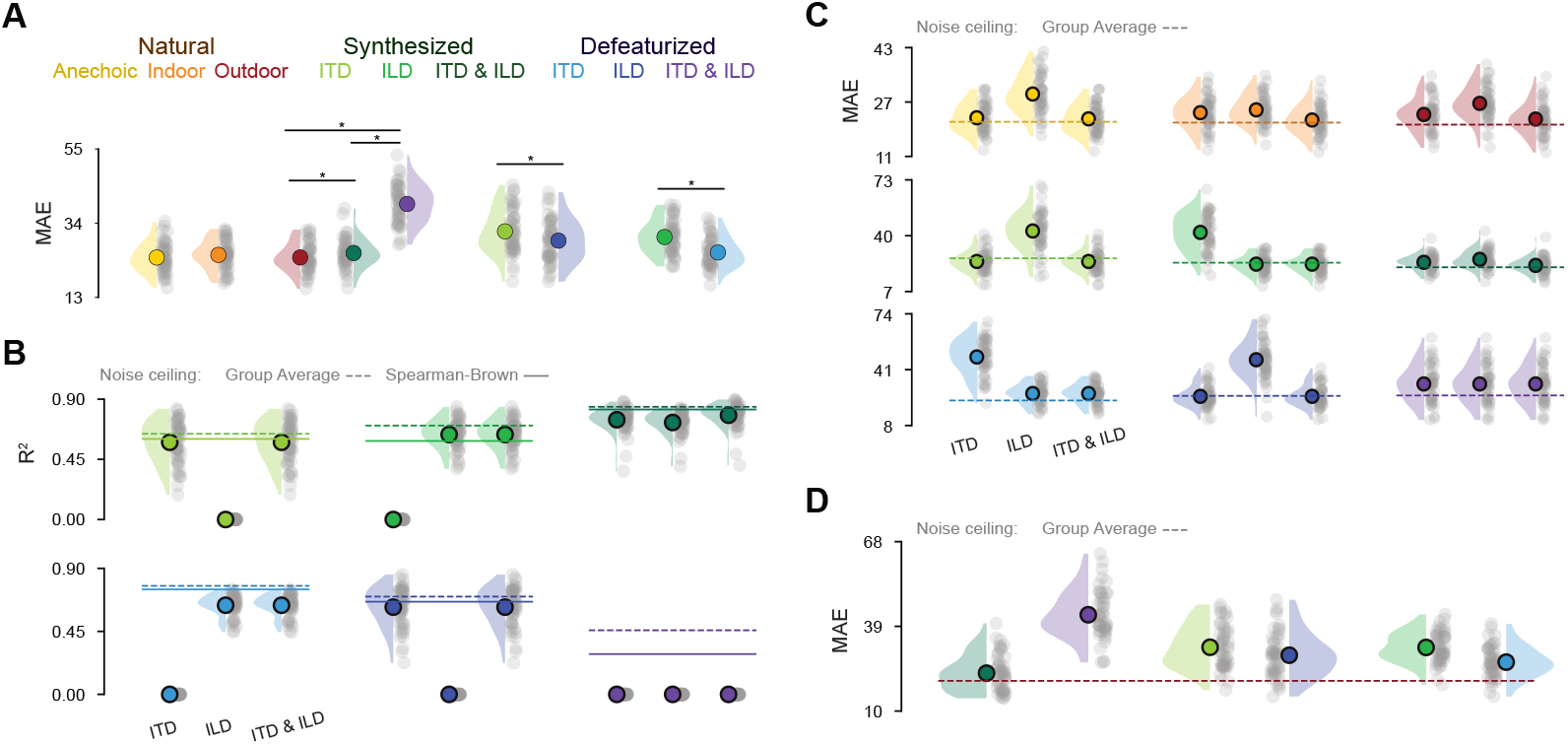
The role of ITD, ILD and third features in shaping human spatial hearing. **a**. Raincloud plots showing the average MAE score in degrees across all azimuth positions in all conditions. Each dot represents a participant, while the circle overlapping the density plot represent the mean. Asterisks signal statistical significance. **b**. Raincloud plots show R^2^ scores for linear regression models fitted to predict human responses on synthesis and defeaturize stimuli using as regressor either the ITD, the ILD or both computed on these sounds. Dots represent participants, while circles indicate the group average. Horizontal lines represent noise ceiling estimates, either from the group average (dotted) or with the Spearman-Brown method (solid). **c**. Raincloud plots show MAE scores from the same linear regression models as in **b** and in Fig. 3A. Horizontal lines represent noise ceiling estimates from the group average. **d**. Raincloud plots show MAE scores for linear regression models predicting human responses to natural outdoor stimuli from responses to synthesis and defeaturize stimuli (same as in Fig. 3B). Horizontal lines represent noise ceiling estimates from the group average in the natural outdoor stimuli.

**Figure S5.**
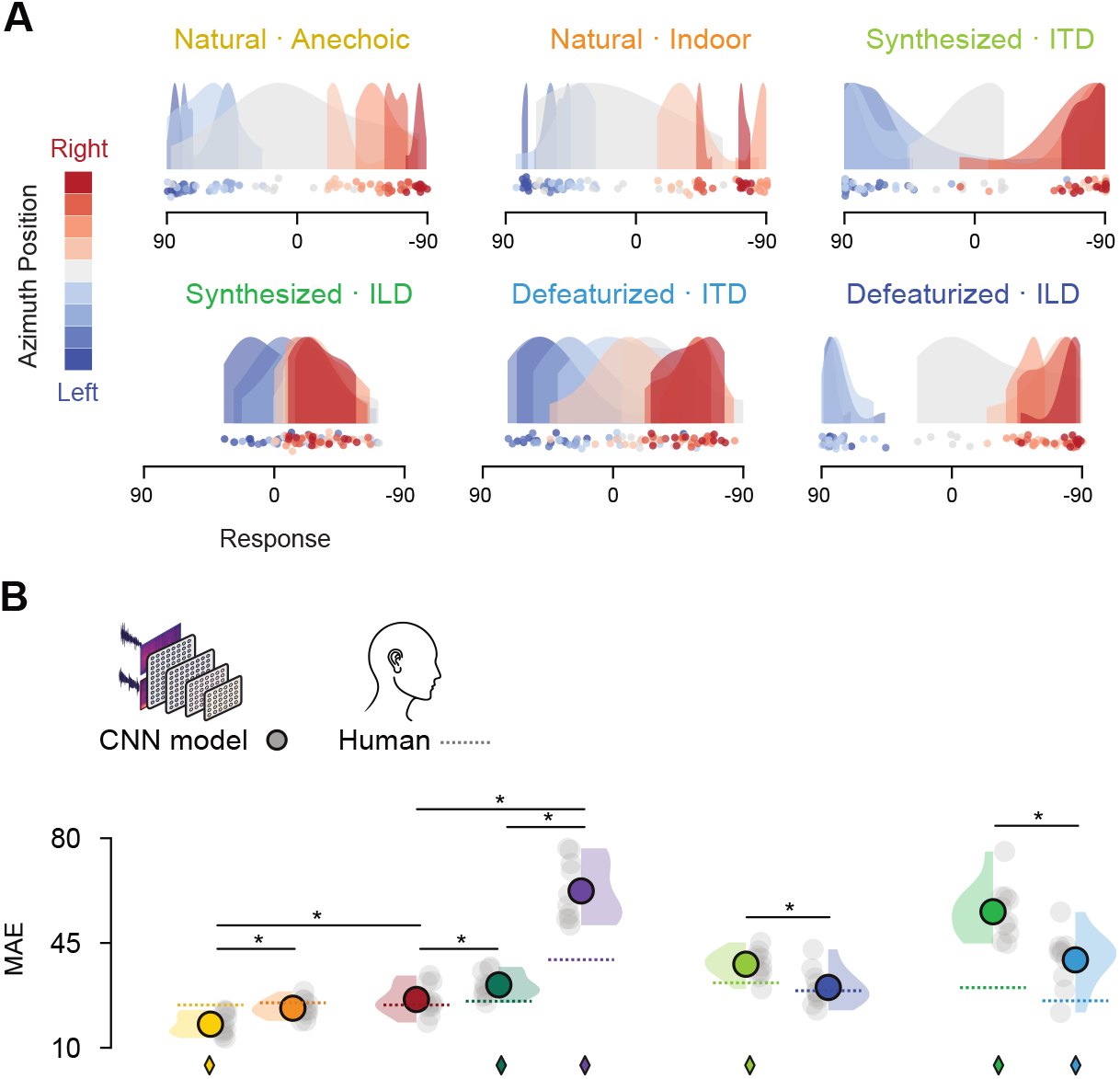
Sound localization performance of deep neural network models. **a**. CNN model response distributions for the natural, synthesis and defeaturize stimuli that were not shown in Fig. 5C. Each dot represents the average response of each model instance to a specific location, color-coded by the azimuth position as expressed by the colorbar. **b**. Raincloud plots showing the average MAE score in degrees across all azimuth positions in all conditions for humans and CNN models. Each dot represents human participants or model instances, while the circle overlapping the density plot represent the mean. Asterisks signal statistical significance.

**Figure S6.**
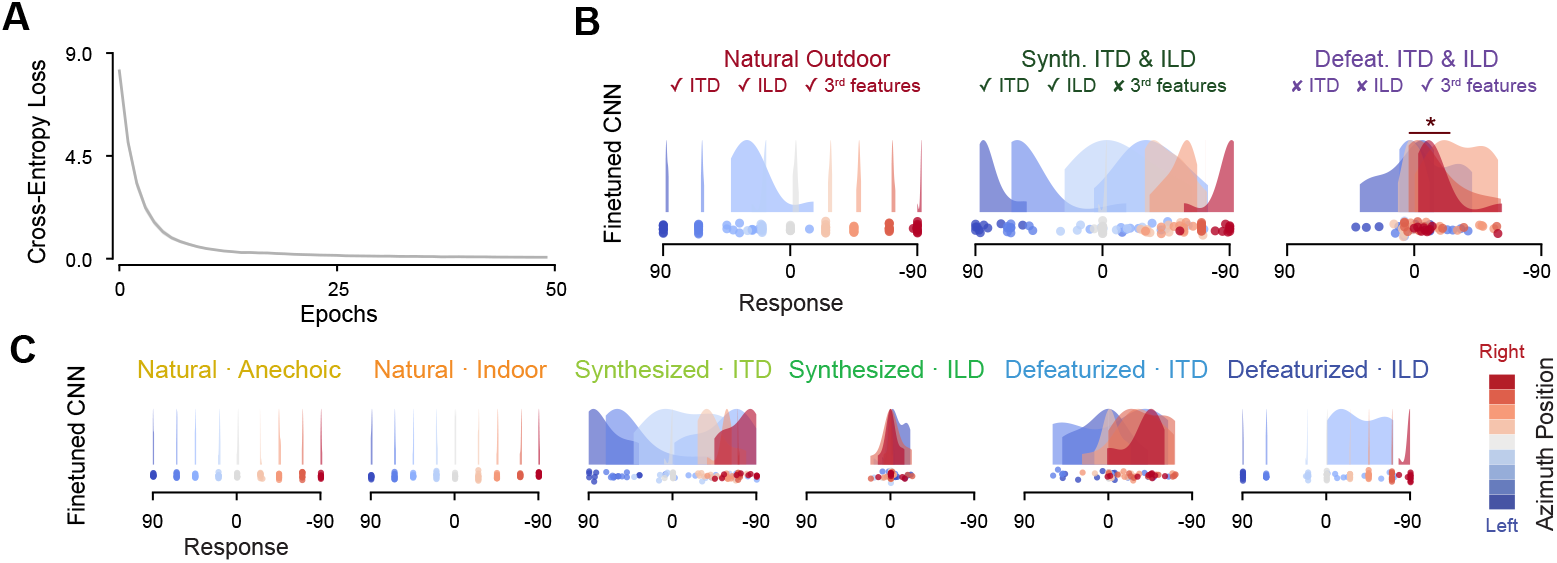
Sound localization performance of finetuned CNN models. **a**. Lineplot showing the loss function across epochs for the finetuning of the CNN models pretrained in the physics simulator. **b**. Model prediction distributions for the natural outdoor, synthesis ITD-ILD and defeaturize ITD-ILD stimuli. Each dot represents the average response of each model to a specific location, color-coded by the azimuth position as expressed by the colorbar. Asterisks represent statistical significance. **c**. Finetuned CNN model response distributions for the natural, synthesis and defeaturize stimuli that were not shown in **b**. (in this layout for consistency with the format of the Sim-trained CNN results presentation). Each dot represents the average response of each model instance to a specific location, color-coded by the azimuth position as expressed by the colorbar.

**Figure S7.**
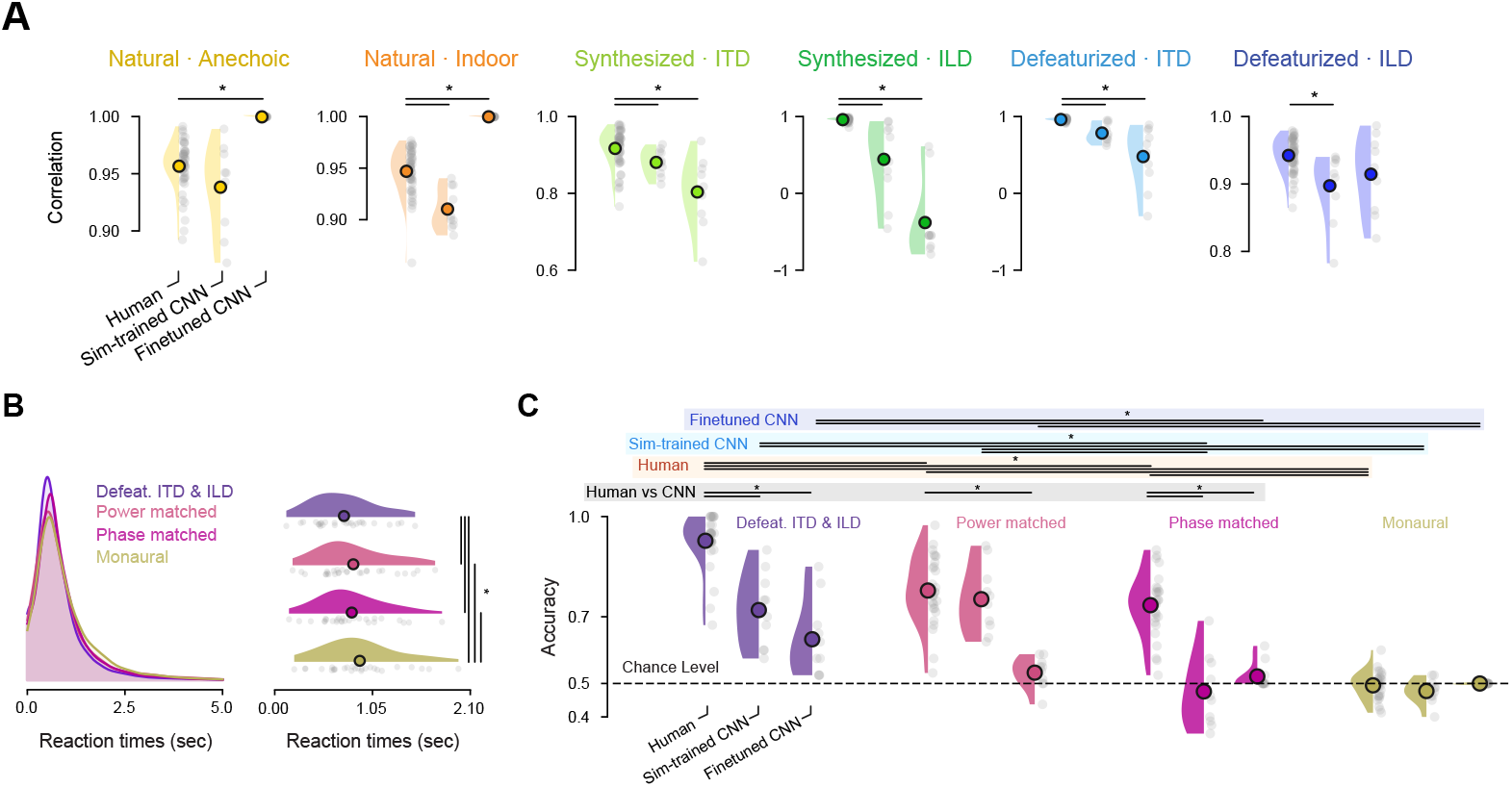
Comparing sound localization in humans and deep neural network models. **a**. Raincloud plots showing Pearson correlation values between the ideal performance and human or model responses. Each dot represents individual human participants or CNN model instances, while the circle overlapping the density plot represent the average across humans or models. Asterisks signal statistical significance, generally as P < 0.05. **b**. Left, density plot showing the reaction time distribution across the four stimulus types we presented to human listeners. Right, raincloud plots showing the average reaction time per human participant, represented by each grey dot. Asterisks indicate statistical significance. **c**. Raincloud plots showing the accuracy of humans or CNN models in the 2AFC sound localization task across stimulus types. For power, phase and monaural matched stimuli, the accuracy is assessed as an average from both sides (e.g., for stimuli located to the left we considered the accuracy as the average from the generate stimuli with the left and right channels as reference). Each dot represents individual human participants or CNN model instances, while the circle overlapping the density plot represent the average across humans or models. Horizontal dotted black line indicates chance level performance. Asterisks indicate statistical significance, color-coded by whether the comparisons were tested within human or model performance or between human and models.

## References

1. van der Heijden, K., Rauschecker, J. P., de Gelder, B. & Formisano, E. Cortical mechanisms of spatial hearing. Nat. Rev. Neurosci. 20, 609–623 (2019).

2. Blauert, J. Spatial Hearing: The Psychophysics of Human Sound Localization. (MIT press, 1997).

3. Hofman, P. M. Relearning sound localization with new ears. Nat. Neurosci. 1, (1998).

4. Rayleigh, Lord. XII. On our perception of sound direction. Lond. Edinb. Dublin Philos. Mag. J. Sci. 13, 214–232 (1907).

5. Batteau, D. W. The role of the pinna in human localization. Proc. R. Soc. Lond. B Biol. Sci. 168, 158–180 (1967).

6. Grothe, B., Pecka, M. & McAlpine, D. Mechanisms of Sound Localization in Mammals. Physiol. Rev. 90, 983–1012 (2010).

7. Butler, R. A. & Flannery, R. The spatial attributes of stimulus frequency and their role in monaural localization of sound in the horizontal plane. Percept. Psychophys. 28, 449–457 (1980).

8. Middlebrooks, J. C. & Green, D. M. Sound Localization by Human Listeners. Annu. Rev. Psychol. 42, 135–159 (1991).

9. Macpherson, E. A. & Middlebrooks, J. C. Listener weighting of cues for lateral angle: The duplex theory of sound localization revisiteda). J Acoust Soc Am 111, (2002).

10. Krizhevsky, A., Sutskever, I. & Hinton, G. E. ImageNet Classification with Deep Convolutional Neural Networks. in Advances in Neural Information Processing Systems vol. 25 (Curran Associates, Inc., 2012).

11. LeCun, Y., Bengio, Y. & Hinton, G. Deep learning. Nature 521, 436–444 (2015).

12. Richards, B. A. et al. A deep learning framework for neuroscience. Nat. Neurosci. 22, 1761–1770 (2019).

13. Saxe, A., Nelli, S. & Summerfield, C. If deep learning is the answer, what is the question? Nat. Rev. Neurosci. 22, 55–67 (2021).

14. Zador, A. et al. Catalyzing next-generation Artificial Intelligence through NeuroAI. Nat. Commun. 14, 1597 (2023).

15. Doerig, A. et al. The neuroconnectionist research programme. Nat. Rev. Neurosci. 24, 431–450 (2023).

16. Hinton, G. et al. Deep neural networks for acoustic modeling in speech recognition: The shared views of four research groups. IEEE Signal Process. Mag. 29, 82–97 (2012).

17. Mnih, V. et al. Human-level control through deep reinforcement learning. Nature 518, 529–533 (2015).

18. Kaufmann, E. et al. Champion-level drone racing using deep reinforcement learning. Nature 620, 982–987 (2023).

19. Yamins, D. L. K. et al. Performance-optimized hierarchical models predict neural responses in higher visual cortex. Proc. Natl. Acad. Sci. 111, 8619–8624 (2014).

20. Cadieu, C. F. et al. Deep neural networks rival the representation of primate IT cortex for core visual object recognition. PLoS Comput. Biol. 10, e1003963 (2014).

21. Yamins, D. L. & DiCarlo, J. J. Using goal-driven deep learning models to understand sensory cortex. Nat. Neurosci. 19, 356–365 (2016).

22. Kell, A. J. E., Yamins, D. L. K., Shook, E. N., Norman-Haignere, S. V. & McDermott, J. H. A Task-Optimized Neural Network Replicates Human Auditory Behavior, Predicts Brain Responses, and Reveals a Cortical Processing Hierarchy. Neuron 98, 630–644.e16 (2018).

23. Kell, A. J. & McDermott, J. H. Deep neural network models of sensory systems: windows onto the role of task constraints. Curr. Opin. Neurobiol. 55, 121–132 (2019).

24. Kietzmann, T. C. et al. Recurrence is required to capture the representational dynamics of the human visual system. Proc. Natl. Acad. Sci. 116, 21854–21863 (2019).

25. Zhuang, C. et al. Unsupervised neural network models of the ventral visual stream. Proc. Natl. Acad. Sci. 118, e2014196118 (2021).

26. Macpherson, T. et al. Natural and Artificial Intelligence: A brief introduction to the interplay between AI and neuroscience research. Neural Netw. 144, 603–613 (2021).

27. Feather, J., Leclerc, G., Mądry, A. & McDermott, J. H. Model metamers reveal divergent invariances between biological and artificial neural networks. Nat. Neurosci. 26, 2017–2034 (2023).

28. Van Der Heijden, K. & Mehrkanoon, S. Goal-driven, neurobiological-inspired convolutional neural network models of human spatial hearing. Neurocomputing 470, 432–442 (2022).

29. Francl, A. & McDermott, J. H. Deep neural network models of sound localization reveal how perception is adapted to real-world environments. Nat. Hum. Behav. 6, 111–133 (2022).

30. Stecker, G. C. & Hafter, E. R. Temporal weighting in sound localization. J. Acoust. Soc. Am. 112, 1046–1057 (2002).

31. Kawashima, T. & Sato, T. Perceptual limits in a simulated “Cocktail party”. Atten. Percept. Psychophys. 77, 2108–2120 (2015).

32. Norman-Haignere, S., Kanwisher, N. G. & McDermott, J. H. Distinct Cortical Pathways for Music and Speech Revealed by Hypothesis-Free Voxel Decomposition. Neuron 88, 1281–1296 (2015).

33. Plakal, M. & Ellis, D. YAMNet https://github.com/tensorflow/models/tree/master/research/audioset/yamnet. (2019).

34. Howard, A. G. et al. MobileNets: Efficient Convolutional Neural Networks for Mobile Vision Applications. Preprint at 10.48550/arXiv.1704.04861 (2017).

35. McDermott, J. H. & Simoncelli, E. P. Sound Texture Perception via Statistics of the Auditory Periphery: Evidence from Sound Synthesis. Neuron 71, 926–940 (2011).

36. Lindsay, G. W. & Bau, D. Testing methods of neural systems understanding. Cogn. Syst. Res. 82, 101156 (2023).

37. Vilas, M. G., Adolfi, F., Poeppel, D. & Roig, G. Position: an inner interpretability framework for AI inspired by lessons from cognitive neuroscience. in Proceedings of the 41st International Conference on Machine Learning vol. 235 49506–49522 (JMLR.org, Vienna, Austria, 2024).

38. Ju, H., Juan, R., Gomez, R., Nakamura, K. & Li, G. Transferring policy of deep reinforcement learning from simulation to reality for robotics. Nat. Mach. Intell. 4, 1077–1087 (2022).

39. Cai, T., Rakerd, B. & Hartmann, W. M. Computing interaural differences through finite element modeling of idealized human heads. J. Acoust. Soc. Am. 138, 1549–1560 (2015).

40. Hartmann, W. M., Rakerd, B., Crawford, Z. D. & Zhang, P. X. Transaural experiments and a revised duplex theory for the localization of low-frequency tones. J. Acoust. Soc. Am. 139, 968–985 (2016).

41. Thurlow, W. R. & Runge, P. S. Effect of induced head movements on localization of direction of sounds. J. Acoust. Soc. Am. 42, 480–488 (1967).

42. Greco, A. & Siegel, M. A spatiotemporal style transfer algorithm for dynamic visual stimulus generation. Nat. Comput. Sci. (2024) doi:10.1038/s43588-024-00746-w.

43. Dürschmid, S. et al. Hierarchy of prediction errors for auditory events in human temporal and frontal cortex. Proc. Natl. Acad. Sci. 113, 6755–6760 (2016).

44. Jiang, X., Han, C., Li, Y. A. & Mesgarani, N. Exploring Self-Supervised Contrastive Learning of Spatial Sound Event Representation. Preprint at 10.48550/arXiv.2309.15938 (2023).

45. Fei, Z., Fan, M. & Huang, J. A-JEPA: Joint-Embedding Predictive Architecture Can Listen. Preprint at 10.48550/arXiv.2311.15830 (2024).

46. Greco, A., Moser, J., Preissl, H. & Siegel, M. Predictive learning shapes the representational geometry of the human brain. Nat. Commun. 15, 9670 (2024).

47. Greco, A., Rastelli, C., Bonetti, L., Braun, C. & Caria, A. Neural signatures of predictive coding underlying the acquisition of incidental sensory associations. 2025.05.16.654429 Preprint at 10.1101/2025.05.16.654429 (2025).

48. Greco, A., Rastelli, C., Ubaldi, A. & Riva, G. Immersive exposure to simulated visual hallucinations modulates high-level human cognition. Conscious. Cogn. 128, 103808 (2025).

49. Bonetti, L. et al. Shared and modality-specific brain networks underlying predictive coding of temporal sequences. 2025.07.16.665207 Preprint at 10.1101/2025.07.16.665207 (2025).

50. Mingard, C., Rees, H., Valle-Pérez, G. & Louis, A. A. Deep neural networks have an inbuilt Occam’s razor. Nat. Commun. 16, 220 (2025).

51. Huh, M. et al. The Low-Rank Simplicity Bias in Deep Networks. Preprint at 10.48550/arXiv.2103.10427 (2023).

52. De Palma, G., Kiani, B. & Lloyd, S. Random deep neural networks are biased towards simple functions. in Advances in Neural Information Processing Systems vol. 32 (Curran Associates, Inc., 2019).

53. Nakkiran, P. et al. SGD on Neural Networks Learns Functions of Increasing Complexity. Preprint at 10.48550/arXiv.1905.11604 (2019).

54. Gunasekar, S., Lee, J. D., Soudry, D. & Srebro, N. Implicit Bias of Gradient Descent on Linear Convolutional Networks. in Advances in Neural Information Processing Systems vol. 31 (Curran Associates, Inc., 2018).

55. Soudry, D., Hoffer, E., Nacson, M. S., Gunasekar, S. & Srebro, N. The Implicit Bias of Gradient Descent on Separable Data. J. Mach. Learn. Res. 19, 1–57 (2018).

56. Valle-Perez, G., Camargo, C. Q. & Louis, A. A. Deep learning generalizes because the parameter-function map is biased towards simple functions. in International Conference on Learning Representations (2018).

57. Arpit, D. et al. A Closer Look at Memorization in Deep Networks. in Proceedings of the 34th International Conference on Machine Learning 233–242 (PMLR, 2017).

58. Jo, J. & Bengio, Y. Measuring the tendency of CNNs to Learn Surface Statistical Regularities. Preprint at 10.48550/arXiv.1711.11561 (2017).

59. Shah, H., Tamuly, K., Raghunathan, A., Jain, P. & Netrapalli, P. The Pitfalls of Simplicity Bias in Neural Networks. in Advances in Neural Information Processing Systems vol. 33 9573–9585 (Curran Associates, Inc., 2020).

60. Vasudeva, B., Shahabi, K. & Sharan, V. Mitigating Simplicity Bias in Deep Learning for Improved OOD Generalization and Robustness. Trans. Mach. Learn. Res. (2024).

61. Schnupp, J. W. H. & Carr, C. E. On hearing with more than one ear: lessons from evolution. Nat. Neurosci. 12, 692–697 (2009).

62. Jin, C., Corderoy, A., Carlile, S. & van Schaik, A. Contrasting monaural and interaural spectral cues for human sound localization. J. Acoust. Soc. Am. 115, 3124–3141 (2004).

63. Grumiaux, P.-A., Kitić, S., Girin, L. & Guérin, A. A survey of sound source localization with deep learning methods. J. Acoust. Soc. Am. 152, 107–151 (2022).

64. Morgado, P., Li, Y. & Nvasconcelos, N. Learning Representations from Audio-Visual Spatial Alignment. in Advances in Neural Information Processing Systems vol. 33 4733–4744 (Curran Associates, Inc., 2020).

65. Marentakis, G. Spatial Audio for Multimodal Location Monitoring. Interact. Comput. 33, 564–582 (2021).

66. Sun, P. et al. Both Ears Wide Open: Towards Language-Driven Spatial Audio Generation. in International Conference on Learning Representations (2024).

67. Rascon, C. & Meza, I. Localization of sound sources in robotics: A review. Robot. Auton. Syst. 96, 184–210 (2017).

68. Cho, H. et al. Auptimize: Optimal Placement of Spatial Audio Cues for Extended Reality. in Proceedings of the 37th Annual ACM Symposium on User Interface Software and Technology 1–14 (ACM, Pittsburgh PA USA, 2024). doi:10.1145/3654777.3676424.

69. Corrêa De Almeida, G., Costa de Souza, V., Da Silveira Júnior, L. G. & Veronez, M. R. Spatial Audio in Virtual Reality: A systematic review. in Proceedings of the 25th Symposium on Virtual and Augmented Reality 264–268 (Association for Computing Machinery, New York, NY, USA, 2024). doi:10.1145/3625008.3625042.

70. Luntz, M. et al. Sound localization in patients with cochlear implant—preliminary results. Int. J. Pediatr. Otorhinolaryngol. 64, 1–7 (2002).

71. Dorman, M. F. et al. Interaural Level Differences and Sound Source Localization for Bilateral Cochlear Implant Patients. Ear Hear. 35, 633–640 (2014).

72. Veugen, L. C. E. et al. Horizontal sound localization in cochlear implant users with a contralateral hearing aid. Hear. Res. 336, 72–82 (2016).

73. Körtje, M., Baumann, U., Stöver, T. & Weissgerber, T. Sensitivity to interaural time differences and localization accuracy in cochlear implant users with combined electric-acoustic stimulation. PLOS ONE 15, e0241015 (2020).

74. Schäfer, E. et al. Activities of the Right Temporo-Parieto-Occipital Junction Reflect Spatial Hearing Ability in Cochlear Implant Users. Front. Neurosci. 15, (2021).

75. Peirce, J. W. PsychoPy—psychophysics software in Python. J. Neurosci. Methods 162, 8–13 (2007).

76. Shinn-Cunningham, B. G., Desloge, J. G. & Kopco, N. Empirical and modeled acoustic transfer functions in a simple room: effects of distance and direction. in Proceedings of the 2001 IEEE Workshop on the Applications of Signal Processing to Audio and Acoustics (Cat. No.01TH8575) 183–186 (2001). doi:10.1109/ASPAA.2001.969573.

77. Allen, J. B. & Berkley, D. A. Image method for efficiently simulating small-room acoustics. J. Acoust. Soc. Am. 65, 943–950 (1979).

78. Gaik, W. Combined evaluation of interaural time and intensity differences: Psychoacoustic results and computer modeling. J. Acoust. Soc. Am. 94, 98–110 (1993).

79. Chung, W., Carlile, S. & Leong, P. A performance adequate computational model for auditory localization. J. Acoust. Soc. Am. 107, 432–445 (2000).

80. May, T., van de Par, S. & Kohlrausch, A. A Probabilistic Model for Robust Localization Based on a Binaural Auditory Front-End. IEEE Trans. Audio Speech Lang. Process. 19, 1–13 (2011).

81. Jiang, S., Wu, L., Yuan, P., Sun, Y. & Liu, H. Deep and CNN fusion method for binaural sound source localisation. J. Eng. 2020, 511–516 (2020).

82. Glorot, X., Bordes, A. & Bengio, Y. Deep Sparse Rectifier Neural Networks. in Proceedings of the Fourteenth International Conference on Artificial Intelligence and Statistics 315–323 (JMLR Workshop and Conference Proceedings, 2011).

83. Ioffe, S. & Szegedy, C. Batch normalization: Accelerating deep network training by reducing internal covariate shift. in International conference on machine learning 448–456 (PMLR, 2015).

84. Srivastava, N., Hinton, G., Krizhevsky, A., Sutskever, I. & Salakhutdinov, R. Dropout: a simple way to prevent neural networks from overfitting. J. Mach. Learn. Res. 15, 1929–1958 (2014).

85. Kingma, D. P. & Ba, J. Adam: A Method for Stochastic Optimization. in ICLR (eds Bengio, Y. & LeCun, Y.) (2015).

86. Spearman, C. Correlation calculated from faulty data. Br. J. Psychol. 3, 271 (1910).

87. Brown, W. Some experimental results in the correlation of mental abilities 1. Br. J. Psychol. 1904-1920 3, 296–322 (1910).

88. Benjamini, Y. & Hochberg, Y. Controlling the False Discovery Rate: A Practical and Powerful Approach to Multiple Testing. J. R. Stat. Soc. Ser. B Methodol. 57, 289–300 (1995).

